# Engineered nanosponges mitigate peripheral stress-induced neuroinflammation and restore cognitive function

**DOI:** 10.64898/2026.09.16.752213

**Authors:** Sajeeshkumar Madhurakkat Perikamana, Biswanath Maity, Rokas Dargis, Raina Kikani, Hilal Ahmad Rather, Paris Brown, Giwon cho, Hunter Newman, Lavonia Duncan, Gaurav Arya, Shyni Varghese

## Abstract

Systemic inflammation is increasingly recognized as a key driver of neuroinflammation and cognitive dysfunction, particularly in the aging population. Yet, therapeutic interventions that broadly attenuate circulating inflammatory mediators without suppressing host immunity remain limited. Here, we report the development of taurine-functionalized hyaluronic acid nanosponges (HA-Tau) that blunt systemic inflammatory cascades to protect against downstream neurocognitive impairment. Using molecular docking calculations and experimental validations, we show that taurine functionalization enhances multivalent interactions with diverse cytokines, enabling broad-spectrum sequestration of inflammatory proteins from both murine and human plasma while preserving the intrinsic hypochlorous acid neutralizing ability of taurine. In aged mice undergoing orthopedic surgery, systemic administration of HA-Tau nanosponges lowered the levels of circulating inflammatory mediators, preserved blood–brain barrier integrity, and attenuated glial cell activation. These effects were accompanied by improved hippocampal neuronal activity and spatial working memory in mice. By dampening peripheral inflammatory surges, the nanosponges limit peripheral-to-central inflammatory signaling without directly targeting the central nervous system. Collectively, these findings demonstrate systemic inflammatory modulation could be an effective strategy for mitigating peripheral insult-induced neuroinflammation and cognitive decline, and position HA-Tau nanosponges as a versatile biomaterial platform for treating inflammation-driven disorders.

## Introduction

Neuroinflammation is a central pathological process underlying a wide range of acute and chronic neurological disorders [1, 2]. Although the initial activation of resident glial cells, particularly astrocytes and microglia, provides a protective role, chronic or dysregulated neuroinflammatory cascades—characterized by sustained, elevated levels of cytokines, chemokines, and reactive oxygen species—disrupts neural homeostasis, impair cognitive function, and contributes to the progression of neurodegenerative conditions [3]. Importantly, neuroinflammation is not solely initiated by direct injury to the central nervous system (CNS); peripheral inflammatory insults are increasingly recognized as potent system-level drivers, particularly in aging populations [4]. Systemic infections, autoimmune disorders, trauma, and peripheral injuries such as orthopaedic surgery are well-documented triggers of neuroinflammation [5-8]. The widespread neurological sequelae observed following SARS-CoV-2 infection, including persistent neurocognitive impairment and “brain fog” further highlights the impact of peripheral inflammation on CNS homeostasis and function [6]. Interestingly, even sterile insults such as cardiac and orthopedic surgeries can elicit robust neuroinflammatory responses resulting in postoperative cognitive dysfunction in elderly patients [9-11]. Consequently, modulating peripheral immune activation offers a highly accessible therapeutic approach for mitigating central pathologies.

Peripheral inflammatory insults trigger systemic surges of circulating mediators, such as interleukin-6 (IL-6), interleukin-1β (IL-1β), and tumor necrosis factor-α (TNF-α). Under homeostatic conditions, the blood–brain barrier (BBB)—a highly specialized endothelial interface stabilized by tight junctions—regulates the entry of inflammatory peripheral mediators into the CNS [12]. However, sustained systemic inflammation compromises BBB integrity, increasing vascular permeability and facilitating the infiltration of circulating cytokines, damage-associated molecular patterns, and peripheral immune cells into the brain parenchyma [13]. This infiltration fuels the activation of resident glial populations and amplify neuroinflammatory signaling, thereby linking peripheral immune dysregulation to CNS pathology [4, 13].

Current therapeutic strategies for managing systemic inflammation predominantly rely on broad immunosuppressive drugs, such as corticosteroids, or agents targeting individual cytokines [14, 15]. Although effective in acute, localized contexts, these interventions are often associated with adverse effects, including systemic immunosuppression and increased susceptibility to secondary infections [16, 17]. Furthermore, monoclonal antibodies targeting specific cytokines often exhibit limited efficacy due to redundant and complex signaling networks governing systemic inflammation. Complete inhibition of cytokine signaling may impair physiological tissue repair and essential host defense mechanisms [18]. These limitations highlight a critical need for new innovative approaches capable of attenuating excessive, multi-cytokine inflammatory signaling without inducing profound systemic immunosuppression.

To this end, we developed a biomaterial-based decoy approach designed to broadly modulate systemic inflammation and thereby prevent the trans-signaling events that drive neuroinflammation. We engineered nanosponges composed of hyaluronic acid (HA) functionalized with taurine (HA-Tau), an endogenous amino-sulfonic acid with well-established anti-inflammatory and antioxidant properties [19, 20]. While taurine directly scavenges hypochlorous acid to mitigate oxidative stress-associated tissue damage, we reasoned that its incorporation to an HA backbone would synergistically enhance the physical sequestration of circulating inflammatory mediators. Indeed, amino acid functionalization of HA improves its capacity to bind cytokines and growth factors [21], and sulfonate-bearing biomaterials exhibit broad electrostatically driven affinity for diverse cytokines and proteins [22, 23]. We therefore hypothesized that HA–Tau nanosponges could dampen systemic inflammatory surges by simultaneously sequestering multiple circulating inflammatory mediators, thereby preserving BBB integrity and halting the neuroinflammatory cascade triggered by peripheral insults (Fig. 1). To test this, we characterized the ability of HA-Tau nanosponges to sequester plasma-resident biomolecules in both mouse and human plasma. We then demonstrated that the nanosponge-mediated modulation of systemic inflammation preserves BBB integrity, attenuates neuroinflammation, and preserves cognitive function in an aged mouse model of orthopedic surgery-induced neurocognitive impairment.

**Fig. 1.**
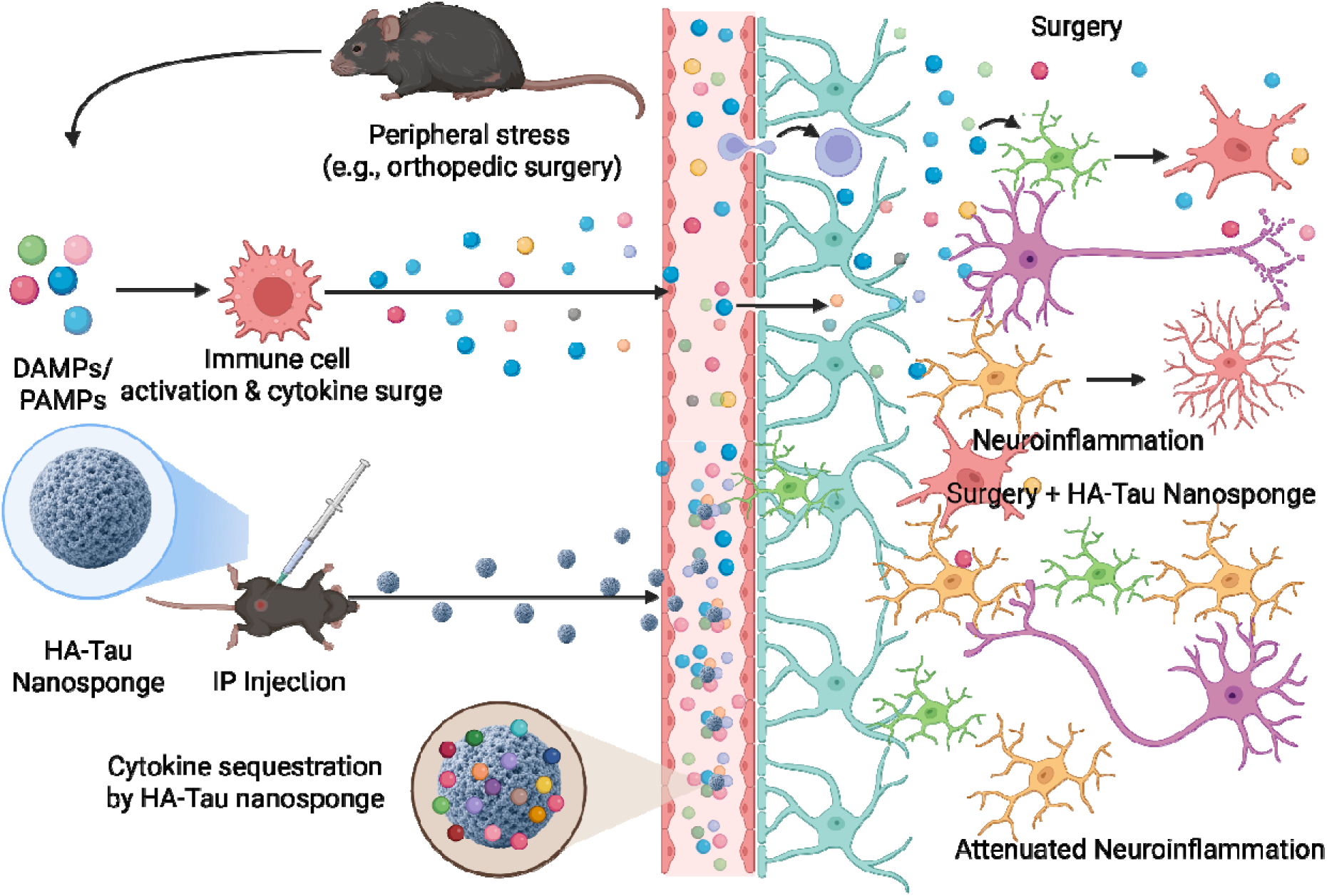
Schematic illustration of the proposed mechanism by which HA–Tau nanosponge sequester multiple circulating inflammatory mediators following peripheral injury, thereby preserving BBB integrity and limiting the subsequent neuroinflammatory cascade. The figure was created using Biorender.com.

## Results

### Computational assessment of taurine functionalization on cytokine sequestration

While the capacity of taurine to neutralize hypochlorous acid is well established, whether taurine-functionalized biomaterials can actively sequester pro-inflammatory cytokines remains elusive. To address this, we first performed *in silico* molecular docking calculations to determine whether functionalizing HA with taurine (HA-Tau) enhances its binding affinity for key pro-inflammatory cytokines. Specifically, we evaluated the binding poses of HA and HA-Tau molecules with several key inflammatory cytokines and calculated the corresponding binding energies. We selected mouse IL-6, IL-1β, and TNF, as well as human IL-6 for this analysis due to the established role of these cytokines in driving inflammaging and systemic inflammation-induced cognitive impairment [24, 25].

To minimize computational complexity and avoid confounding interactions arising from multiple binding sites along long polymer chains, we used a minimal HA trimer base model composed entirely of wild-type units (HA_NNN_). Taurine (T) was then systematically conjugated to monomeric units within the trimer to generate a library of functionalized HA oligomers with distinct substitution patterns: HA_TNN, HA_NTN, HA_NNT, HA_TTN, HA_TNT, HA_NTT, and HA_TTT (Supplementary Fig.1 A-E). Because HA monomers are chiral, sequence permutations (e.g., HA_TNN versus HA_NNT) yield non-superimposable stereoisomers that may exhibit distinct binding behaviors. Hence, we evaluated all possible substitution patterns to systematically distinguish the effects of taurine substitution ratio from those of its spatial arrangement along the HA trimer on cytokine binding.

Molecular docking was performed using AutoDock Vina.[26] To identify distinct binding pockets, the predicted binding poses were grouped using average-linkage hierarchical clustering based on root-mean-square deviation (RMSD)[27], with a clustering cutoff of 15 Å (Supplementary Fig. 2). The number of identified clusters varied across the target cytokines, with predicted docking energies ranging from approximately −10 to −5 kcal/mol (Fig. 2A, B and Supplementary Fig. 3). Across all examined cytokines, the lowest-energy binding pose consistently corresponded to a taurine-substituted HA trimer. In addition, taurine-functionalized HA trimers showed binding pockets that were not accessible to unsubstituted HA (Fig. 2C). Although these additional binding modes were generally less thermodynamically favorable, as indicated by their weaker predicted binding energies, their emergence suggests that taurine incorporation broadens the accessible interaction landscape of HA. Such an expanded binding repertoire may promote cytokine sequestration through multivalent and sequence-dependent interactions, consistent with prior reports demonstrating that multivalent polymer-protein interactions can enhance apparent binding affinity and broaden target engagement [28].

**Fig. 2.**
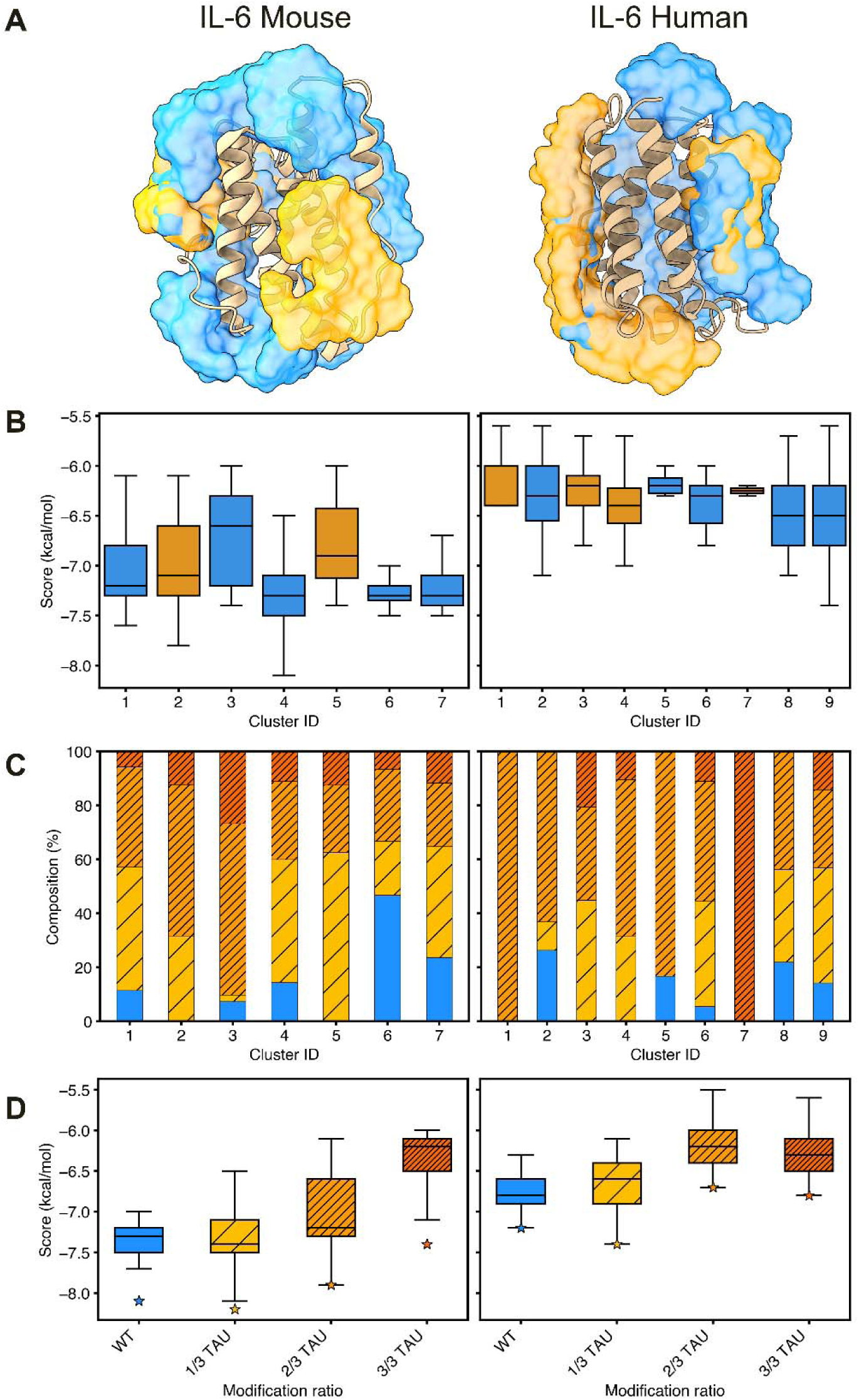
HA trimer docking results onto mouse (left) and human (right) IL-6. A) Clusters visualized onto each mouse and human IL-6, colored based on if the cluster contains only unsubstituted HA (orange) or includes Tau-substituted HA (blue). B) Predicted docking scores grouped by cluster, again colored based on the cluster’s composition. C) Composition of each cluster: blue is unsubstituted HA (blue), 1/3 HA-Tau (yellow), 2/3 HA-Tau (orange), and fully substituted (dark orange). D) Predicted docking scores grouped by HA-Tau ratio, following the same color scheme as panel C. Stars indicate the best predicted binding score.

Analysis of predicted binding scores as a function of taurine substitution revealed a non-monotonic relationship between cytokine binding affinity and the degree of taurine substitution (Fig. 2D and Supplementary Fig. 4). Trimers with a single taurine substitution (HA_TNN_, HA_NTN_, HA_NNT_) showed more favorable binding energies than unsubstituted HA (HA_NNN_). In contrast, further increasing the degree of substitution progressively reduced binding affinity, with doubly substituted trimers (HA_TTN, HA_TNT, and HA_NTT) displaying less favorable docking scores, and fully substituted trimers (HA_TTT) displaying the weakest scores across all examined cytokines. Collectively, these findings suggest that partial taurine substitution may enhance cytokine sequestration by strengthening interactions at binding pockets already accessible to unsubstituted HA while also enabling access to additional binding pockets not sampled by unsubstituted HA. Conversely, excessive or complete taurine substitution may disrupt favorable polymer-cytokine interactions, thereby diminishing overall binding affinity.

### Fabrication of HA–Tau nanosponges and sequestration of plasma-resident biomolecules

Hyaluronic acid (HA; Mw 40 kDa) was functionalized with taurine *via* EDC/NHS-mediated coupling between the carboxyl groups of HA and the primary amine of taurine (Fig. 3A). Based on our promising docking results, we synthesized, by varying the taurine to HA feed ratio, HA derivatives with two levels of substitution: HA–Tau-20 and HA–Tau-40, corresponding to ∼24.83 ± 1.9% and 43.08 ± 2.8% taurine substitution per HA disaccharide repeat unit, respectively. Successful conjugation was confirmed by ^1^H NMR spectroscopy, which revealed characteristic taurine methylene proton resonances at 2.80–2.82 ppm (Supplementary Fig. 5A and B). HA-Tau molecules with substitution levels above 40% were not used, as excessive substitution could reduce their functional efficacy.

**Fig. 3.**
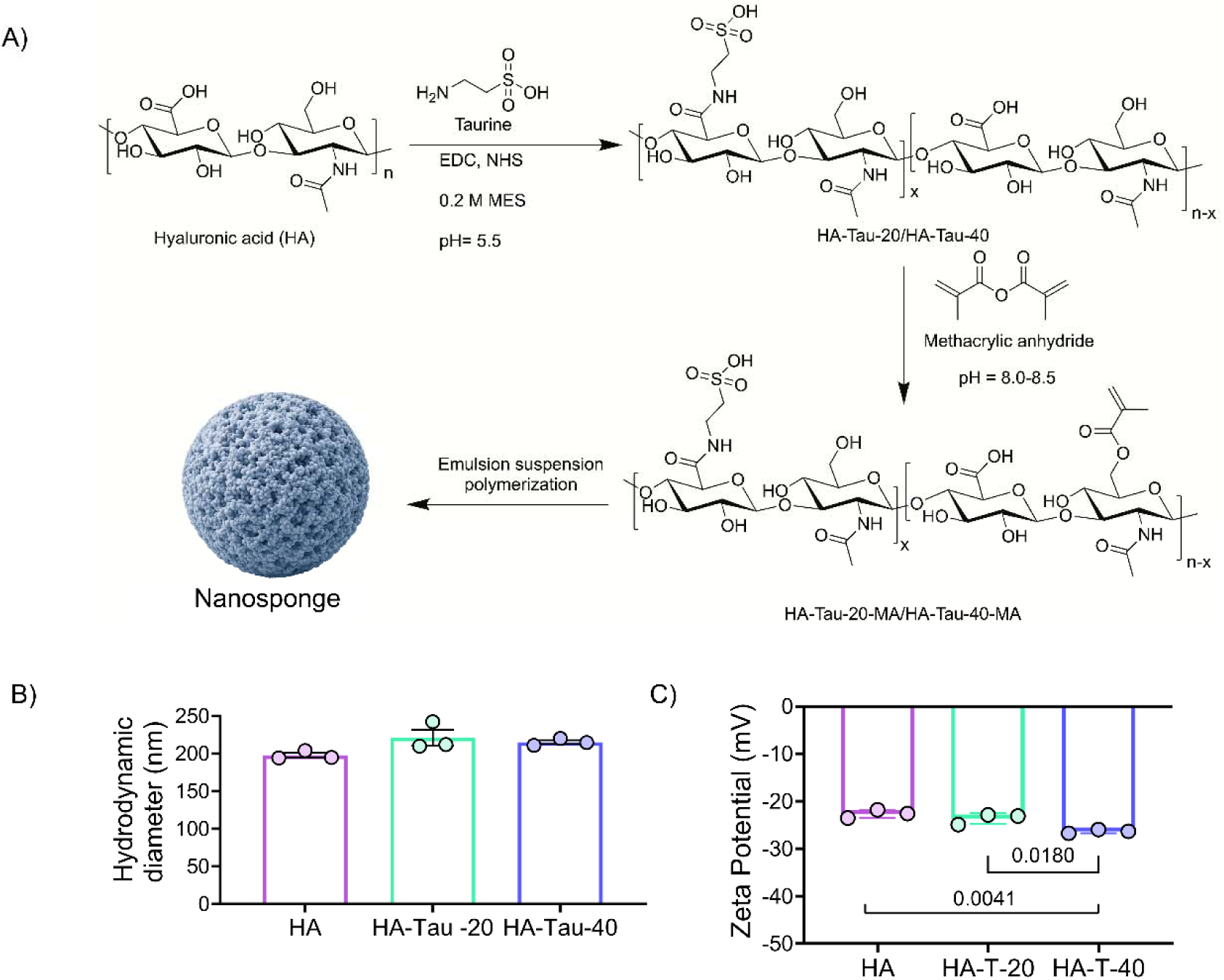
Synthesis and characterization of taurine-modified HA nanosponges. A) Schematic showing the multistep chemical conjugation of taurine and methacrylate groups to hyaluronic acid (HA), followed by emulsion suspension polymerization to form the nanosponges (n=3). B) Hydrodynamic diameter of the HA and taurine-modified HA nanosponges. (C) Surface charge of the HA and taurine-modified HA nanosponges (n=3). Data represent mean ± s.e.m. Statistical analyses were performed using one-way ANOVA with Tukey’s multiple comparisons test in GraphPad Prism 10.2.1. P values less than 0.05 are shown.

The HA-Tau molecules were subsequently methacrylated to generate photocrosslinkable HA-Tau-20-MA and HA-Tau-40-MA polymers. Successful methacrylation was confirmed by ^1^H-NMR spectroscopy, which showed characteristic olefinic peaks at 5.75 and 6.2 ppm (Supplementary Fig. 6A and B), and indicated a methacrylation degree of ∼ 27 ± 1.5% per HA repeat unit. The methacrylated polymers were then processed by emulsion suspension polymerization to form the nanosponges. HA nanosponges lacking taurine served as a control. Detailed synthesis and characterization procedures are described in the Methods. Dynamic light scattering (DLS) analysis showed that the HA, HA-Tau-20, and HA-Tau-40 nanosponges possessed mean hydrodynamic diameters of 197.73±5.51 nm, 221.40±18.13 nm, and 215.16±4.57 nm, respectively (Fig. 3B). The corresponding zeta potentials were –30.3 ± 3.4 mV (HA), –34.7 ± 2.8 mV (HA-Tau-20), and –37.9 ± 0.6 mV (HA-Tau-40), showing an increase in negative surface charge with increasing taurine incorporation (Fig. 3C).

We next experimentally assessed whether taurine functionalization promotes protein-sequestration capacity of the HA nanosponges by using two highly abundant plasma-resident proteins, albumin and fibrinogen. HA-Tau nanosponges with varying taurine densities were incubated with 10 µg/mL of bovine serum albumin (BSA) or bovine fibrinogen for 6 h. Quantification of the unbound fraction revealed significantly higher protein sequestration by HA–Tau-40 for both fibrinogen and BSA compared to HA and HA-Tau-20 nanosponges (Supplementary Fig. 7 A and B). Based on these results, HA–Tau-40 nanosponges were used in subsequent experiments unless otherwise specified.

To determine the broad spectrum of plasma proteins captured by HA–Tau nanosponges, we incubated HA-Tau-40 with plasma collected from 22-month-old mice following tibial fracture surgery, a model previously shown to induce systemic inflammation and elevated levels of senescence associated secretory phenotypes (SASP) [29]. Bound proteins were isolated and characterized by LC–MS/MS–based proteomics (Fig. 4A). Proteomic profiling identified ∼548 proteins associated with the nanosponges (Supplementary file 1). Ingenuity Pathway Analysis (IPA) revealed enrichment of pathways associated with systemic inflammation including acute phase response signaling, IL-12 signaling and production in macrophages, the complement cascade, and the coagulation system (Fig. 4B, Supplementary file 2). These findings indicate that HA-Tau nanosponges effectively sequester major mediators of systemic inflammatory cascades. To identify the key upstream regulators of the captured inflammatory proteome, we performed upstream regulator analysis of the proteins involved in the acute phase response signaling pathway (*P* < 0.05; Supplementary file 3). Among the predicted upstream regulators, the pro-inflammatory cytokines IL-6, IL-1β, and TNF-α exhibited higher activation scores and collectively regulated a large subset of sequestered proteins (Fig. 4C). Network analysis further showed that, IL-6/IL-1β/TNF-α–NF-κB-associated inflammatory axis, linking multiple captured acute-phase proteins to inflammatory signaling pathways (Fig. 4G, Supplementary file 3).

**Fig. 4.**
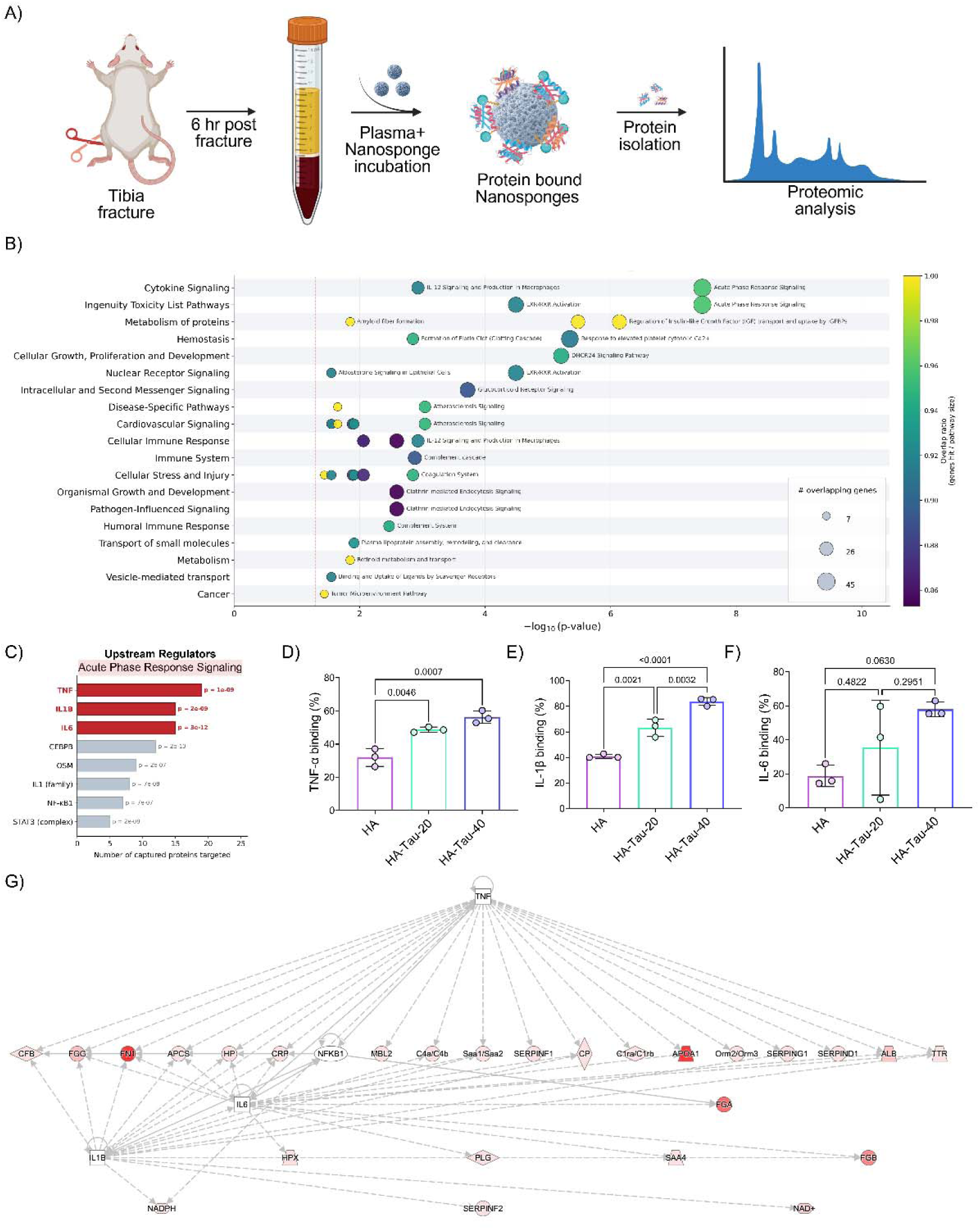
HA–Tau nanosponges sequester circulating inflammatory factors from plasma. A) Schematic of the *ex vivo* plasma proteomics workflow. Plasma collected from 22-month-old mice 6 h after tibial fracture surgery was incubated with HA–Tau-40 nanosponges, followed by isolation of nanosponge-bound proteins and LC–MS/MS-based proteomic analysis (n=1). B) Top enriched pathways identified by Ingenuity Pathway Analysis (IPA) of proteins captured by HA– Tau-40 nanosponges from plasma. The x-axis indicates statistical significance (−log10(P value)), and bubble size represents the number of overlapping proteins. C) Upstream regulator analysis of the proteins associated with acute-phase response signaling among the HA–Tau-40-bound plasma proteome. Top predicted inflammatory regulators are shown. D–F), Quantification of TNF-α (D), IL-1β (E) and IL-6 (F) captured by HA, HA–Tau-20 and HA–Tau-40 nanosponges, as determined by targeted ELISA (n=3). G) Network analysis showing the predicted interactions among inflammatory regulators and proteins captured by HA–Tau-40 nanosponges. Nodes represent proteins or predicted upstream regulators, and connecting lines indicate predicted regulatory or molecular relationships. Data represent mean ± s.e.m. Statistical analyses were performed using one-way ANOVA with Tukey’s multiple comparisons test in GraphPad Prism 10.2.1.

Guided by these proteomic findings, we next used targeted ELISAs to validate cytokine sequestration by HA–Tau nanosponges and to determine the effect of taurine substitution density on cytokine sequestration. HA–Tau nanosponges successfully captured TNF-α, IL-1β, and IL-6 from plasma with HA–Tau-40 exhibiting the highest sequestration efficiency (Fig. 4D–F). Collectively, these results demonstrate that taurine functionalization substantially enhances the ability of HA nanosponges to sequester key pro-inflammatory cytokines.

In addition to cytokine sequestration, we evaluated whether HA–Tau nanosponges retained the intrinsic HOCl-neutralizing activity of taurine [20]. Primary human neutrophils exposed to post-fracture plasma from aged mice induced elevated HOCl production; however, treatment with HA–Tau nanosponges significantly reduced HOCl levels in the culture supernatant (Supplementary Fig. 7C). Together, these findings demonstrate that HA–Tau nanosponges simultaneously sequester broad spectrum cytokines and neutralize reactive oxygen species, thereby providing a multifaceted approach to modulate systemic inflammation.

### *In Vivo* biodistribution and biocompatibility of HA-Tau Nanosponges

To evaluate the biodistribution of HA-Tau nanosponges, we administered Cy5.5-labeled nanosponges intraperitoneally (i.p) into aged mice. Longitudinal analyses of blood samples showed detectable fluorescence at 30 minutes post-injection, which peaked at 6 h before gradually declining (Supplementary Fig. 8A). Twenty-four h post-injection, mice were euthanized and major organs, including the liver, spleen, heart, kidneys, lungs, and brain, were harvested and imaged using an IVIS imaging system. HA-Tau nanosponges were accumulated predominantly in the liver with substantially weaker fluorescence signals detected in other organs (Supplementary Fig. 8B). Given the substantial hepatic accumulation, hematoxylin and eosin (H&E) staining of liver sections was performed to assess acute tissue toxicity. Histological examination revealed no evidence of acute liver toxicity or structural abnormalities associated with HA-Tau nanosponge administration (Supplementary Fig. 8C). To further evaluate cytocompatibility, primary mouse macrophages were incubated with either HA or HA-Tau-40 nanosponges for 24 h. Live/dead staining showed that neither formulation induced detectable cytotoxicity compared with untreated controls (Supplementary Fig. 8D).

To determine whether HA-Tau nanosponges crossed the BBB and entered the brain parenchyma, brain sections were stained for CD31, an endothelial cell marker, and the spatial distribution of Cy5.5 fluorescence relative to the vascular network was assessed. Fluorescence imaging revealed that Cy5.5 signals were predominantly localized within CD31-positive blood vessels, with minimal fluorescence extending beyond the vascular endothelium. These results indicate that HA–Tau nanosponges remained largely confined within the cerebral vasculature and exhibited minimal extravasation into the brain parenchyma (Supplementary Fig. 8E).

### HA-Tau nanosponge-mediated dampening of systemic inflammation preserves BBB integrity

To determine whether nanosponge-mediated sequestration of inflammatory mediators could mitigate peripheral stress–induced BBB disruption and subsequent neuroinflammation, we employed a preclinical model of orthopedic surgery. Tibial fracture surgery in aged animals elicits a systemic inflammatory surge that promotes BBB breaching and neuroinflammation, consistent with clinical observations in older adults [29, 30]. Aged mice subjected to tibial fracture received either HA–Tau nanosponges or saline 30 min post-surgery, as described in the Methods section. Plasma samples were collected 6 h post-surgery for downstream analysis, (Fig. 5A)— a timepoint chosen because circulating SASP-associated cytokines peak transiently at 6 h post-fracture [29]. ELISA measurements of plasma for IL-1β, IL-6, and TNF-α showed significantly lower circulating levels in mice that received HA-Tau-nanosponges compared with saline treated controls, consistent with our *ex vivo* findings demonstrating the ability of HA-Tau nanosponges to sequester various plasma-resident inflammatory mediators (Fig. 5B).

**Fig. 5.**
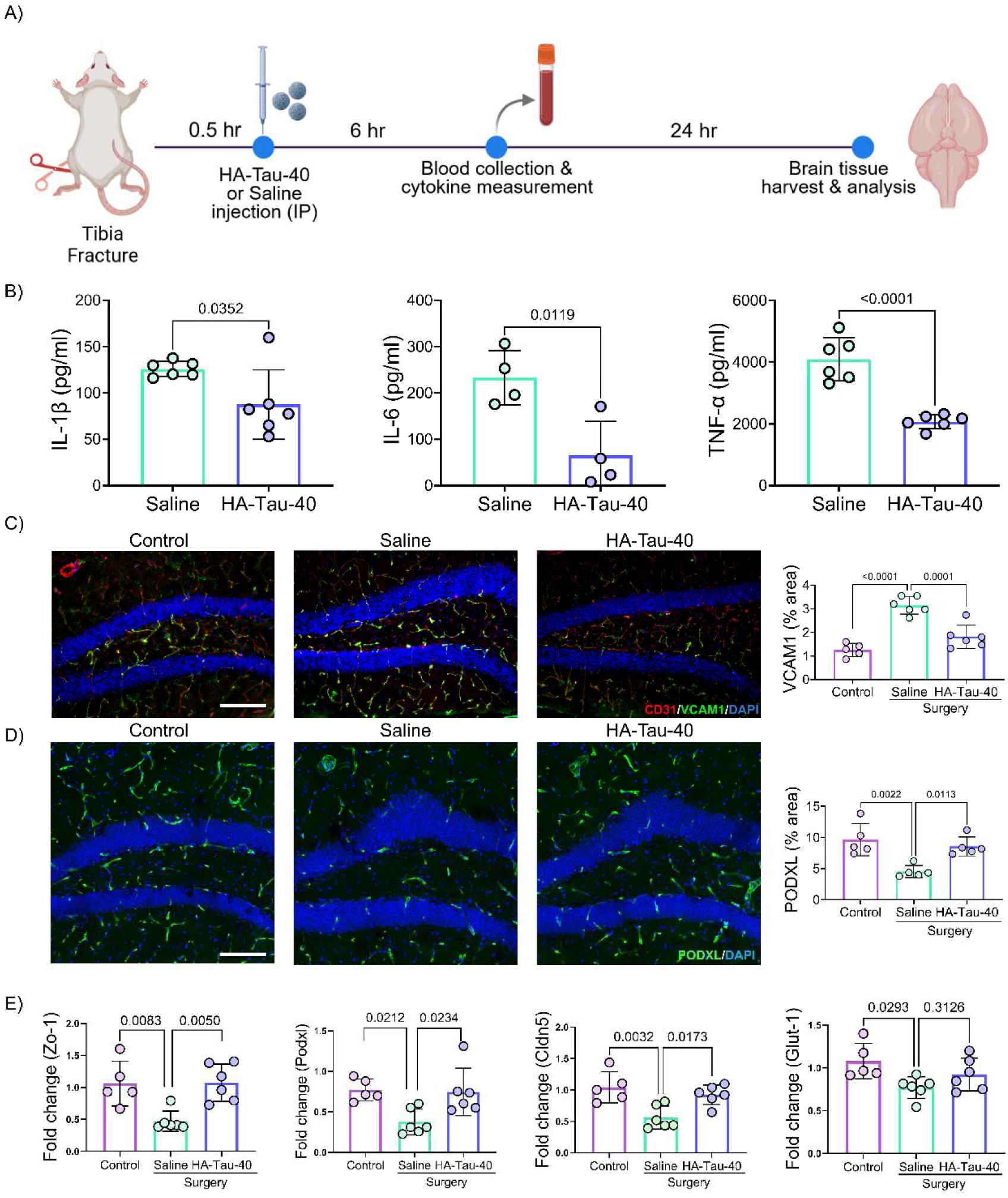
HA–Tau nanosponges attenuate systemic inflammation and preserve blood–brain barrier integrity following tibial fracture. A) Schematic representation of the experimental design with the timeline. Plasma was collected 6 h after surgery for cytokine analysis, and brain tissue was collected 24 h after surgery for assessment of BBB integrity. B) Plasma concentrations of IL-1β, IL-6 and TNF-α measured 6 h after tibial fracture in saline- and HA–Tau-40-treated mice (n=4-6). C) Representative hippocampal images showing VCAM-1 (green), CD31 (red) and DAPI (blue) staining in non-surgery control and saline or HA–Tau-40-treated surgery groups. Scale bar, 100 μm. Quantification of VCAM-1-positive area is shown on the right (n=5-6). D) Representative hippocampal images showing PODXL (green) and DAPI (blue) staining in control and saline or HA–Tau-40-treated fracture groups. Scale bar, 100 μm. Quantification of PODXL-positive area is shown on the right (n=5). E) Relative expression of BBB-associated genes, including *ZO-1*, *PODXL*, *CLDN5* and *GLUT1*, in the hippocampus of control, saline-treated fracture and HA–Tau-40-treated fracture mice, quantified by RT–qPCR (n=5-6). Data represent mean ± s.e.m. Statistical analyses were performed using two-Student’s t-test or one-way ANOVA with Tukey’s multiple comparisons test in GraphPad Prism 10.2.1. P values less than 0.05 are shown.

We next investigated whether attenuating the systemic inflammatory surge could preserve BBB integrity. Endothelial activation and BBB inflammation were assessed by immunostaining for vascular cell adhesion molecule-1 (VCAM-1) [31, 32]. Tibial fracture surgery resulted in a marked increase in VCAM-1 expression of endothelial cells; however, HA–Tau nanosponge treatment substantially attenuated this surgery-induced upregulation (Fig. 5C).

To further characterize BBB structural integrity, we assessed the expression of podocalyxin (PODXL), an endothelial surface glycoprotein implicated in maintenance of vascular barrier function.[33] Aged mice subjected to tibial fracture displayed reduced hippocampal PODXL expression compared to non-injured controls. Administration of HA–Tau nanosponges prevented this decline, maintaining PODXL levels comparable to those observed in no-surgery group (Fig. 5D). Consistent with these findings, RT–qPCR analysis demonstrated that HA-Tau nanosponge treatment attenuated the fracture-induced downregulation of BBB integrity–associated genes, including ZO-1, PODXL, CLDN5, and GLUT-1, in the hippocampus (Fig. 5E).

We further evaluated BBB permeability using Evans Blue extravasation assay. Following systemic dye administration, brains were harvested, fixed, sectioned, and immunostained for CD31 to visualize the vasculature. In saline administered control mice, Evans Blue fluorescence was detected in the perivascular space surrounding multiple blood vessels, indicating BBB leakage. In contrast, in HA–Tau nanosponge–administered mice, the Evans Blue signal remained largely confined to CD31-positive vessels, suggesting reduced vascular permeability and preservation of BBB integrity (Supplementary Fig. 9A and B). Together, these findings demonstrate that the sequestration of systemic inflammatory mediators following peripheral insults attenuates BBB inflammation, maintains BBB integrity, and restricts the transmission of peripheral inflammatory signals to the brain.

### HA -Tau mediated dampening of systemic inflammation attenuated neuroinflammation

We have previously shown that aged animals develop a neuroinflammatory response following tibial fracture surgery as characterized by activation of astrocytes and microglia. Consistent with these findings, immunostaining for the glial markers GFAP and IBA1 revealed a significant increase in astrocyte and microglial activation in the hippocampi of the surgery group relative to non-surgical controls. In contrast, mice administered with HA–Tau nanosponges following surgery showed significantly reduced glial activation, with levels approaching those observed in non-surgical animals (Fig. 6A and B).

**Fig. 6.**
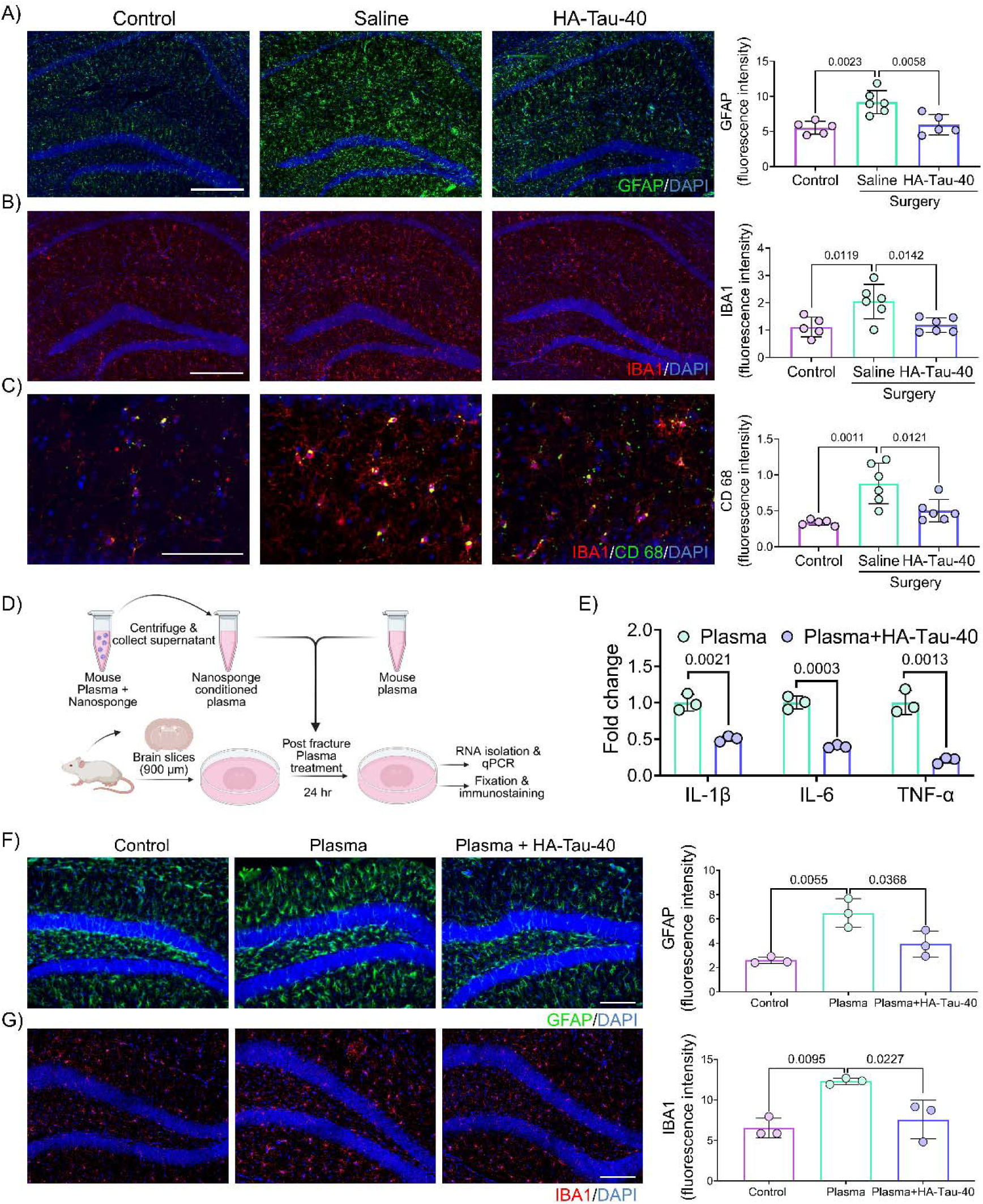
HA–Tau nanosponges attenuate surgery-induced neuroinflammation. A) Representative hippocampal images of GFAP (green) and DAPI (blue) staining and corresponding quantification of GFAP fluorescence intensity in non-surgical control, saline-treated fracture, and HA–Tau-40-treated fracture mice. Scale bar, 100 μm (n=5-6). B) Representative hippocampal images showing IBA1 (red) and DAPI (blue) staining and corresponding quantification of IBA1 fluorescence intensity in the three experimental groups. Scale bar, 100 μm (n=5-6). C) Representative hippocampal images of IBA1 (red), CD68 (green), and DAPI (blue) staining and corresponding quantification of CD68 fluorescence intensity in the three experimental groups. Scale bar, 50 μm (n=5-6). D) Schematic of the *ex vivo* brain explant assay used to assess the effects of circulating inflammatory mediators and their modulation by HA–Tau nanosponges. Plasma collected 6 h after tibial fracture was incubated with HA–Tau-40 nanosponges, and conditioned plasma was subsequently applied to brain explants. E) Relative expression of IL1B, IL6, and TNF in brain explants cultured with fracture plasma or HA–Tau-40-conditioned fracture plasma (n=3). F and G) Representative brain explant images showing GFAP staining following exposure to control medium, fracture plasma, or HA–Tau-40-conditioned fracture plasma. Quantification of GFAP and IBA1 fluorescence intensity is shown on the right. Scale bar, 100 μm (n=3). Statistical analyses were performed using two-tailed Student’s t-test or one-way ANOVA with Tukey’s multiple comparisons test in GraphPad Prism 10.2.1. P values less than 0.05 are shown.

Microglial activation was further examined by immunostaining for CD68, a marker for reactive microglia. Tibial fracture surgery induced a pronounced increase in CD68-positive microglia within the hippocampus, whereas HA–Tau nanosponge treatment substantially attenuated this response, resulting in CD68 expression levels comparable to those of non-surgical controls (Fig. 6C).

To further investigate the contribution of circulating inflammatory mediators, and their modulation by HA-Tau nanosponges, on hippocampal neuroinflammation, we utilized an *ex vivo* brain explant model. Plasma was collected from mice 6 h after tibial fracture surgery and conditioned with HA-Tau nanosponge as described in the Methods (Fig. 6D). Brain explants were cultured in either standard medium, medium supplemented with 5% plasma from mice underwent fracture surgery (unconditioned plasma), or medium supplemented with 5% nanosponge-conditioned plasma. Gene expression analyses after 24 h of culture revealed significant downregulation in the expression of the pro-inflammatory cytokines, TNF-α, IL-6, and IL-1β, in explants exposed to nanosponge-conditioned plasma compared to those cultured in presence of unconditioned plasma (Fig. 6E). Concomitantly, immunostaining for GFAP and IBA1 showed reduced astrocyte and microglial activation in explants exposed to nanosponge conditioned plasma relative to unconditioned plasma controls (Fig. 6F and G).

### HA-Tau nanosponges rescue hippocampal function and spatial working memory from systemic inflammation medicated changes

Given that neuroinflammation can alter neuronal network activity, we next examined whether HA–Tau nanosponge–mediated modulation of systemic inflammation preserved hippocampal function following tibial fracture surgery. Spontaneous neuronal activity was assessed in acute hippocampal explants using a flexible microelectrode array (MEA) platform. Representative raster plots recorded across multiple electrodes revealed higher neuronal spiking activity in hippocampal slices from HA-Tau–nanosponge administered mice compared with saline controls (Fig. 7A). Quantitative analysis showed a significant increase in mean firing frequency in hippocampal tissues from mice that received HA-Tau nanosponges, indicating improved neuronal activity following the sequestration of systemic inflammatory mediators (Fig. 7B). Consistent with these functional improvements, immunofluorescence analysis of hippocampal sections stained for synaptophysin (a presynaptic marker) and PSD-95 (a postsynaptic marker) demonstrated increased colocalization of synaptic puncta in HA-Tau–treated mice relative to saline-treated mice (Fig. 7C). Quantitative analysis confirmed a significant increase in synaptophysin/PSD-95 colocalization in HA–Tau-treated mice compared with saline-treated mice, suggesting that HA–Tau nanosponge treatment may help to improve synaptic organization in the hippocampus following surgery-induced systemic inflammation.

**Fig. 7.**
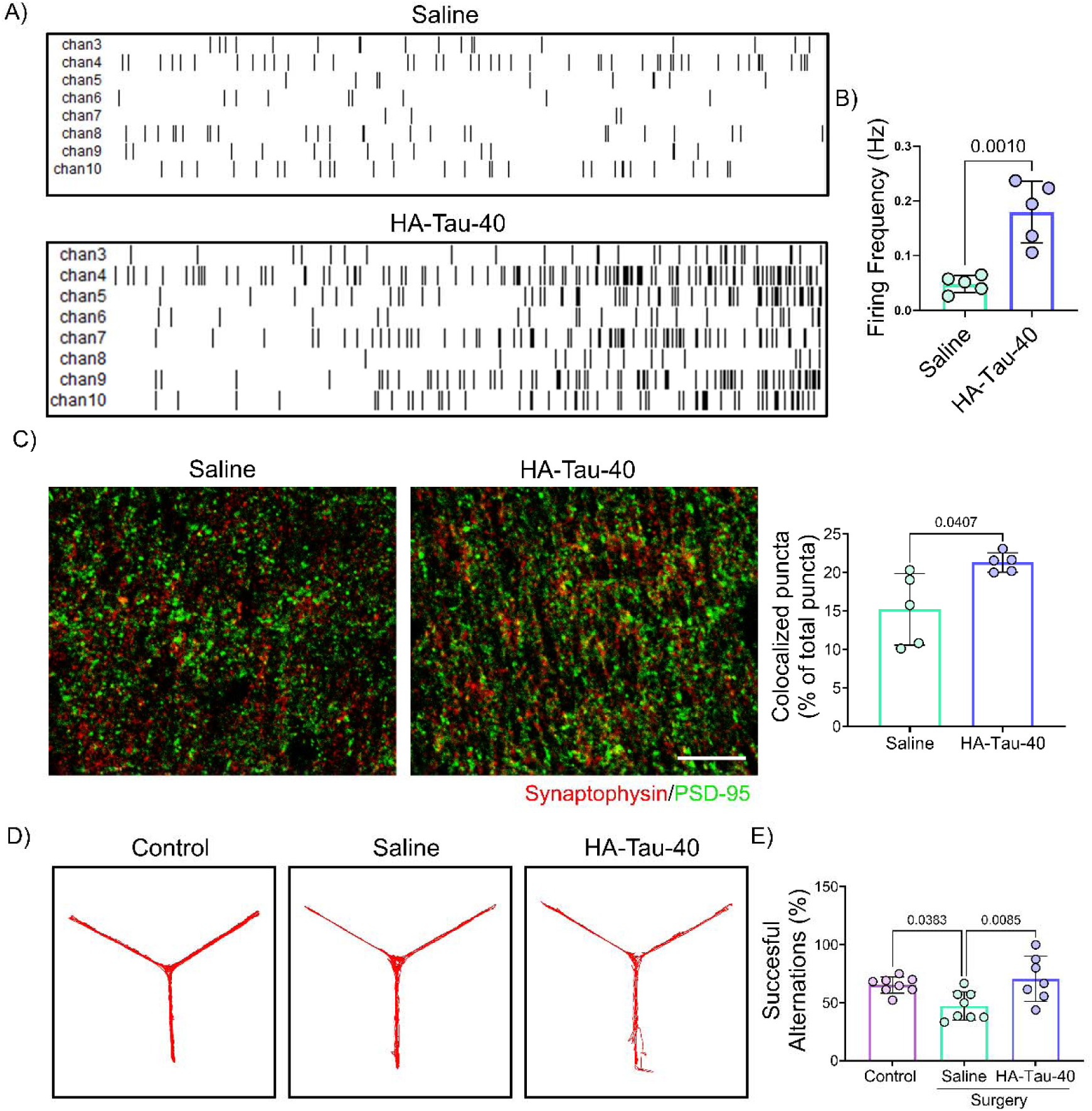
HA–Tau nanosponges preserve hippocampal neuronal activity, synaptic organization and spatial working memory following tibial fracture. A) Representative raster plots of spontaneous neuronal activity recorded from hippocampal slices harvested from saline or HA–Tau-40-treated mice following tibial fracture surgery using a multielectrode array (MEA) platform. B) Quantification of mean neuronal firing frequency in hippocampal slices from saline and HA–Tau-40-treated mice (n=5). C) Representative hippocampal images showing synaptophysin and PSD-95 staining in saline and HA–Tau-40-treated mice. Scale bar, 10 μm. Quantification of synaptophysin/PSD-95 colocalized puncta, expressed as a percentage of total puncta, is shown on the right (n=5). D) Representative Y-maze arm-entry sequences from non-surgical control, saline treated fracture, and HA–Tau-40-treated fracture mice. E) Quantification of spontaneous alternation in the Y maze analysis (n=7-8). Statistical analyses were performed using two-tailed Student’s t-test or one-way ANOVA with Tukey’s multiple comparisons test in GraphPad Prism 10.2.1. P values less than 0.05 are shown.

We next examined whether this reduction in neuroinflammation translated into improved cognitive function. Spatial working memory was assessed using Y-maze spontaneous alternation test, which measures the ability of mice to remember and explore previously unvisited arms [34]. Mice subjected to tibial fracture surgery exhibited significantly reduced spontaneous alternation compared with non-surgical controls, indicative of impaired short-term spatial working memory. Notably, HA–Tau nanosponge administration markedly improved spontaneous alternation performance, restoring it to levels comparable to those observed in non-surgical animals (Fig. 7D and E). Collectively, these findings suggest that systemic administration of HA-Tau nano sponges ameliorates surgery-induced deficits in hippocampal neuronal activity, synaptic integrity, and spatial working memory in aged mice, likely through attenuation of peripheral inflammation and its downstream neuroinflammatory consequences.

### HA-Tau nanosponges sequester inflammatory mediators from aged human plasma and protect BBB in an in vitro model

To determine whether HA–Tau nanosponges similarly sequester circulating inflammatory mediators from human plasma, we incubated the nanosponges with plasma obtained from aged human donors similar to the mouse plasma studies. Aged human plasma was used as previous studies have shown that it induces BBB breaching and promotes neuroinflammation [32]. HA– Tau nanosponges were incubated with human plasma for 6 h, after which the nanosponge-bound proteins were isolated and analyzed by LC–MS/MS–based proteomics. Across three independent donors, proteomic profiling of nanosponge-bound protein corona identified ∼570–590 proteins per plasma sample, with ∼540 proteins shared among all three donors, indicating a highly conserved protein corona composition despite inter-donor variability (Supplementary Fig. 10 A and B, Supplementary file 4). Pathway enrichment analysis showed that the sequestered proteins were predominantly associated with inflammation- and immune-related pathways, including the complement cascade, acute phase response signaling, IL-15 signaling, and the coagulation system (Fig. 8A, Supplementary file 5). Upstream regulator analysis of acute phase response– associated proteins identified the pro-inflammatory cytokines IL-6, IL-1β, and TNF-α among the most significant predicted upstream regulators, collectively targeting a large number of captured proteins, similar to the mouse plasma analysis (Fig. 8B, Supplementary file 6). Network analysis further showed that these cytokines converged on NF-κB, forming a highly interconnected regulatory hub that links the majority of the nanosponge-bound acute phase proteins to inflammatory cascades (Fig. 8D). Targeted ELISA confirmed the sequestration of TNF-α, IL-1β, and IL-6, with taurine substitution substantially enhancing cytokine sequestration compared with unsubstituted HA nanosponges (Fig. 8C).

**Fig. 8.**
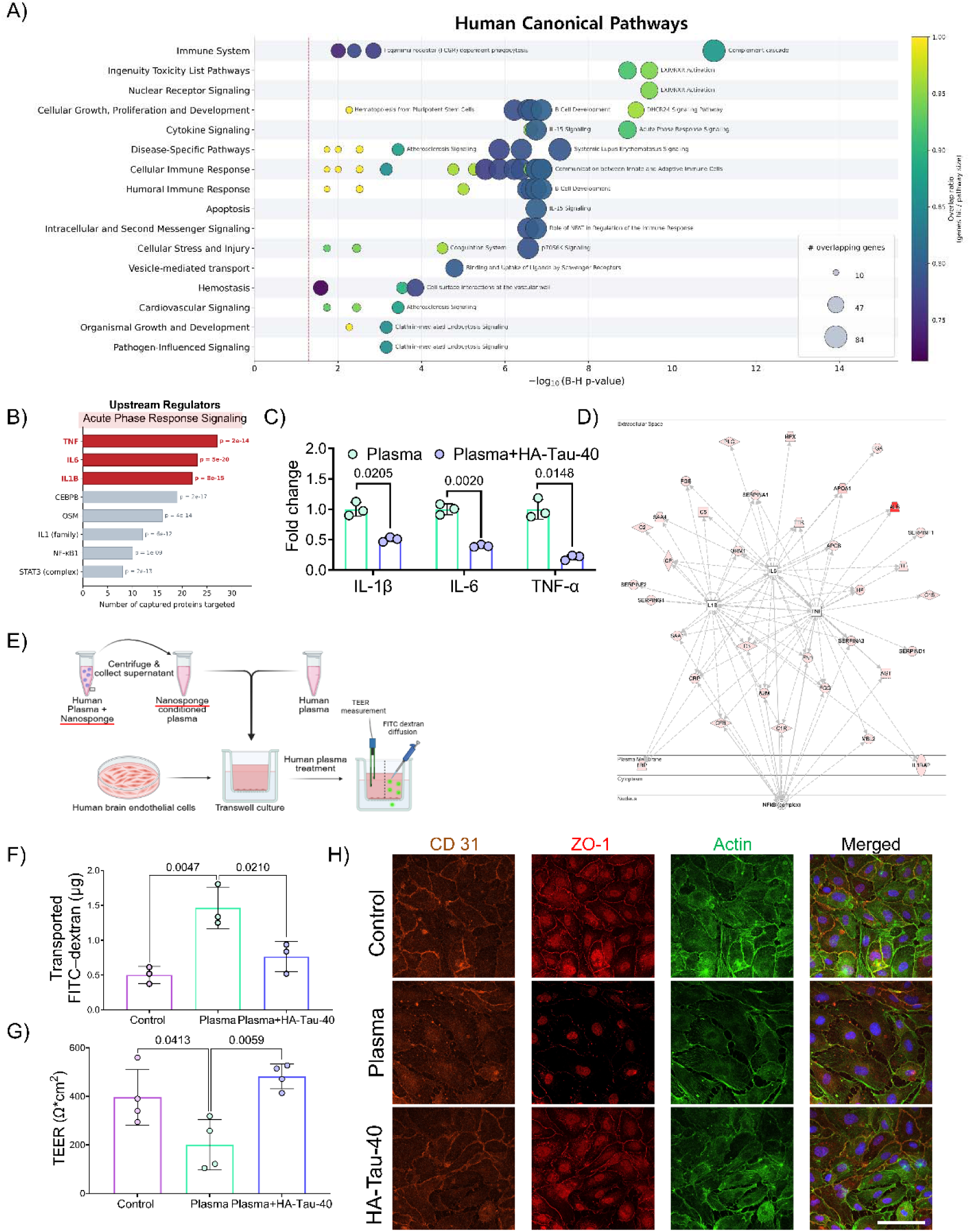
HA–Tau nanosponges sequester inflammatory mediators from aged human plasma and preserve blood–brain barrier function *in vitro*. A) Top enriched pathways identified by pathway analysis of proteins captured by HA–Tau-40 nanosponges from aged human plasma. The x-axis indicates statistical significance (−log10 adjusted *P* value), and bubble size represents the number of overlapping proteins (n=3). B) Upstream regulator analysis of proteins associated with the acute-phase response signaling pathway among the HA–Tau-40-bound human plasma proteome, highlighting the top predicted inflammatory regulators. C) Quantification of IL-1β, IL-6 and TNF-α captured by HA and HA–Tau-40 nanosponges from aged human plasma, as quantified by ELISA (n=3). D) Network analysis showing predicted interactions among inflammatory regulators and proteins associated with the acute-phase response signaling pathway captured by HA–Tau-40 nanosponges. E) Schematic of the *in vitro* human blood–brain barrier model and experimental workflow. Human brain microvascular endothelial cells were exposed to untreated human plasma or plasma conditioned with HA–Tau-40 nanosponges, followed by assessment of barrier function. F) FITC-dextran (10 kDa) transport across the endothelial monolayer following exposure to control medium, aged human plasma, or HA–Tau-40-conditioned plasma (n=3). G) Trans-endothelial electrical resistance (TEER) measurements of endothelial monolayers under different conditions (n=4). H) Representative immunofluorescence images showing CD31 (orange), ZO-1 (red), F-actin (green), and DAPI (blue) in endothelial monolayers exposed to control medium, aged human plasma, or HA–Tau-40-conditioned plasma. Scale bar, 100 μm. Statistical analyses were performed using two-tailed Student’s t-test or one-way ANOVA with Tukey’s multiple comparisons test in GraphPad Prism 10.2.1. P values less than 0.05 are shown.

To evaluate the effect of human plasma following protein sequestration on BBB integrity, we employed an *in vitro* human BBB model consisting of primary human brain microvascular endothelial cells. Barrier function was assessed by measuring the permeability of fluorescein isothiocyanate (FITC)-dextran (10 kDa) and by trans-endothelial electrical resistance (TEER) (Fig. 8D). Exposure to aged human plasma significantly increased FITC-dextran transport across the endothelial layer, indicative of compromised barrier integrity. In contrast, plasma conditioned with HA–Tau nanosponges markedly reduced dextran permeability, approaching levels similar to those of control cultures (Fig. 8E). Consistent with these findings, exposure to aged plasma resulted in a significant reduction in TEER values relative to control cultures, whereas TEER measurements of cultures exposed to nanosponge-conditioned plasma remained comparable to those of the control group (Fig. 8F). Immunostaining for the tight junction protein ZO-1 corroborated these observations, demonstrating preservation of endothelial junctional organization when the (Fig. 8G). Together, these findings suggest that HA–Tau nanosponges effectively sequester pro-inflammatory factors from aged human plasma, thereby contributing to the preservation of BBB integrity.

## Discussion

Peripheral inflammatory responses are increasingly recognized as major drivers of neurovascular dysfunction and cognitive impairment following injury or infections, particularly in aging populations [35-38]. Herein, we developed and evaluated a hyaluronic acid–taurine (HA-Tau) nanosponge platform designed to broadly sequester circulating SASP factors and inflammatory mediators, thereby controlling the peripheral-to-central inflammatory axis. Using a preclinical model of orthopedic trauma-induced neuroinflammation, we demonstrate that HA-Tau nanosponges attenuate systemic cytokine surges, preserve BBB integrity, reduce glial activation, and protect against surgery-induced cognitive decline in aged mice. Together, these findings establish peripheral cytokine modulation as a promising and translationally viable strategy for mitigating secondary neuroinflammatory outcomes.

Surgical procedures, such as orthopedic surgery, are strongly associated with postoperative neurocognitive complications in older adults, including delirium and long-term cognitive impairment [30, 39]. These complications are accompanied by a significant increase in circulating pro-inflammatory mediators. Consistent with these clinical observations, tibial fracture surgery in aged mice elicits a robust systemic inflammatory response characterized by elevated levels of circulating cytokines such as IL-6, TNF-α, and IL-1β. This systemic surge was accompanied by BBB breaching, astrocytic and microglial activation, and cognitive impairment. These findings align with the growing concept that systemic inflammatory mediators contribute directly to CNS vulnerability during aging by destabilizing the neurovascular interface and exacerbating glial reactivity. Targeting systemic inflammation with HA–Tau nanosponges shortly after surgery substantially mitigated these neuropathological responses. This suggests that transient modulation of peripheral inflammatory signaling is sufficient to preserve neurovascular homeostasis and safeguard cognitive function following peripheral insults.

Computational analyses revealed that taurine functionalization introduces interaction motifs that promote multivalent binding with cytokines, thereby expanding the sequestration capacity of the nanosponge platform. These in silico predictions were corroborated experimentally through targeted ELISA measurements. Moreover, proteomic profiling revealed broad-spectrum sequestration of inflammatory mediators associated with inflammatory signaling and innate immune activation pathways. Beyond enhancing protein binding, taurine possesses antioxidant properties, including the ability to neutralize neutrophil derived HOCl. *In vitro* studies demonstrated that HA conjugation preserved the HOCl-scavenging activity of taurine. Together the results suggest that the therapeutic effect of HA–Tau nanosponges likely arise from a combination of cytokine sequestration mediated suppression of inflammatory signaling and attenuation of oxidative inflammatory stress.

Biodistribution studies showed that HA-Tau nanosponges remained largely confined to the vasculature compartment, with minimal accumulation in the brain parenchyma, suggesting negligible BBB penetration. These findings suggest that the neuroprotective effects of the HA-Tau nanosponges are largely mediated through modulation of the systemic inflammatory milieu rather than through direct engagement with the CNS. Indeed, HA–Tau nanosponge reduced the levels of circulating inflammatory mediators implicated in BBB destabilization, which directly correlated with the preservation of BBB integrity following fracture surgery. Attenuation of peripheral inflammation was further associated with diminished astrocytic and microglial activation in the hippocampus, preservation of neuronal network activity, and improved performance in behavioral assays assessing spatial working memory. Collectively, these findings support a model in which peripheral cytokine sequestration interrupts feedforward inflammatory signaling between the systemic circulation and the CNS, thereby limiting neurovascular dysfunction and secondary neuroinflammation after trauma.

The translational potential of this approach is supported by our evaluation using human plasma samples from older adults and *in vitro* human BBB models, in which HA–Tau nanosponges effectively reduced inflammatory mediator burden and preserved barrier integrity. Proteomic analyses of nanosponge-bound protein corona in both mouse and human plasma suggested the emergence of the IL-6/IL-1β/TNFα-NFκB axis as the dominant regulatory signature. This cross-species alignment demonstrates that the nanosponges target a broad spectrum of blood-borne factors in inflammatory network rather than a species-specific pathway, underscoring the potential for clinical translation.

The strategy described in this study differs fundamentally from conventional anti-inflammatory therapies such as corticosteroids or monoclonal antibodies, which often rely on broad immunosuppression or selective blockade of individual cytokines [12–16]. Because post-traumatic inflammation is mediated by interconnected and highly redundant cytokine networks, inhibition of a single cytokine may be insufficient to modulate the broader systemic inflammatory response [40, 41]. In contrast, HA–Tau nanosponges function through a decoy-based sequestration mechanism that dampens hyperinflammation without completely suppressing physiological immune function. This broad-spectrum, yet modulatory mode of action enables simultaneous regulation of multiple inflammatory cascades while minimizing the risks associated with overt immunosuppression.

Although this proof-of-concept study demonstrates the efficacy of biomaterial-assisted broad-spectrum sequestration of inflammatory mediators in preserving BBB integrity and protecting against downstream neuroinflammatory and cognitive deficits without the evidence of any acute organ toxicity, widespread application of this approach requires additional studies. In particular, more comprehensive characterization of the clearance kinetics and long-term safety profiles of HA–Tau nanosponges is needed. This study focused primarily on acute neuroinflammatory and cognitive outcomes; however, whether transient cytokine scavenging confers sustained neuroprotection or alters the trajectory of chronic neuroinflammatory remains to be established. Nevertheless, although the current study focused on postoperative neuroinflammation, the underlying therapeutic principle may be broadly applicable to other systemic inflammatory conditions associated with CNS dysfunction, including sepsis, severe infections, and inflammatory disorders. Defining disease-specific dosing regimens, therapeutic windows, and long-term safety profiles will therefore be essential for future translational development.

Nonetheless, we have developed a biomaterial platform, HA–Tau nanosponges, vascularly confined immunomodulatory approach that mitigates neuroinflammation through the sequestration of broad spectrum peripheral inflammatory mediators. By preserving BBB integrity, attenuating glial activation, and protecting cognitive function without directly targeting the CNS, this approach establishes peripheral immune modulation as a promising strategy for safeguarding the aging brain from systemic inflammatory insults. More broadly, these findings highlight the therapeutic potential of targeting the peripheral inflammatory milieu to prevent neurovascular dysfunction and secondary CNS injury across a diverse spectrum of inflammatory diseases.

## Materials and methods

### Materials

Hyaluronic acid (HA, molecular weight 40 kDa, HA40K-5) was purchased from Lifecore, USA. Methacrylic anhydride (276685), N-hydroxysuccinimide (NHS, 130672), taurine (86329), sodium hydroxide (795429), and mineral oil (M5904) were obtained from Millipore Sigma, USA. 1-Ethyl-3-(3-dimethylaminopropyl) carbodiimide hydrochloride (EDC, D1601) was purchased from TCI. Cyanine 5 amine (430C0) was purchased from Lumiprobe. Dialysis tubing with a molecular weight cutoff of 12-14 kDa (SKU: 132706) was obtained from Repligen. ABIL EM90 surfactant (420095-L-151) was supplied by Universal PreservA Chem Inc. Hexane, acetone, ethanol, and dimethyl sulfoxide (DMSO) were purchased from Millipore-Sigma, with the solvents being ACS or spectroscopic grade. NMR spectra were recorded using a 500 MHz Bruker Avance Neo spectrometer.

### Computational analysis

The wild-type hyaluronic acid trimer structure was created using the GAG builder in Glycam [42], and taurine substitutions were performed using a taurine ligand structure [43] from the Protein Data Bank [44] and ChimeraX [45]. Cytokine structures were also acquired via the Protein Data Bank: mouse IL-6 (PDB ID: 2l3y) [46], human IL-6 (PDB ID: 1il6)[47], mouse IL-1β (PDB ID: 2mib) [48], mouse TNF (PDB ID: 1tnf) [49]. Docking calculations were performed using AutoDock VINA [26] onto the entire protein structure with an exhaustiveness value of 128. For each protein, five docking calculations were run per trimer variant, each producing nine docking predictions, totaling 360 predicted conformations per protein. These conformations were clustered using the average linkage method in SciPy [27] based on their RMSD and a 15 angstrom threshold (Supplementary fig. 2). Any cluster consisting of only one structure was removed from analysis. Structural visualizations were performed using ChimeraX.

### HA-Tau-20 Synthesis

Hyaluronic acid (HA, 400 mg) was dissolved in 200 mM MES buffer (pH 5.5) at 10 mg/mL in a 100 mL round-bottom flask at room temperature (RT). EDC (384 mg; two equiv. relative to HA carboxylate group) was added, and the solution was stirred for 15 minutes. NHS (230 mg; 2 equivalent) was then added, and stirring was continued for an additional 15 minutes to form the NHS ester. Taurine (250 mg; two equiv.) was dissolved in PBS (10 mM, 100 mg/mL) and added to the activated HA solution. The reaction was stirred at RT for 24 h, transferred to a dialysis tube (12-14 kDa MWCO), and dialyzed against DI water for 4 days with water changes twice daily. The solution was transferred into 50 mL centrifuge tubes, snap-frozen in liquid nitrogen, and freeze-dried to obtain HA-Tau-20 powder. The percentage of taurine modification was determined using the ^1^H-NMR.

### HA-Tau-40 Synthesis

Hyaluronic acid (HA, 400 mg) was dissolved in 200 mM MES buffer (pH 5.5) at 10 mg/mL in a 100 mL round-bottom flask at RT. EDC (384 mg; two equiv. relative to HA carboxylate group) was added, and the solution was stirred for 15 minutes. NHS (230 mg; 2 equiv.) was then added, and stirring was continued for an additional 15 minutes to form the NHS ester. Taurine (625.8 mg; 5 equiv.) was dissolved in PBS (10 mM, 100 mg/mL) and added to the activated HA solution. The reaction was stirred at RT for 24 h, transferred to a dialysis tube (12-14 kDa MWCO), and dialyzed against deionized water (DI) for 4 days with water changes twice daily. The solution was transferred into 50 mL centrifuge tubes, snap-frozen in liquid nitrogen, and freeze-dried to obtain HA-Tau-40 powder. The percentage of taurine modification was determined using the ^1^H-NMR.

### HA-MA, HA-Tau-20-MA, and HA-Tau-40-MA Synthesis

HA, HA-Tau-20, or HA-Tau-40 (400 mg) was dissolved in DI water at a concentration of 7.5 mg/mL and transferred to a 250 mL double-neck round-bottom flask. The solution was then cooled on ice. Methacrylic anhydride (2.2 mL; approximately 15 equiv. per HA repeating unit) was added dropwise with vigorous stirring. The pH of the reaction mixture was adjusted to 8.0–9.0 by adding 1 N NaOH in small aliquots (typically 100-200 µL of NaOH solution per addition), ensuring the pH does not exceed 9.5. The reaction proceeded for about 4 h on ice. The reaction mixture was then precipitated by gradually adding it dropwise to ice-cold ethanol-acetone (1:1, v/v) to 5-10 times the volume of the reaction mixture, with stirring. The precipitate was stirred for 20 minutes and allowed to sit for an additional 10 minutes to facilitate settling. The precipitate was collected by centrifugation at 6000 g, washed three times with an ice-cold ethanol-acetone (1:1) mixture, and redissolved in distilled water (DI). The polymer solution was transferred into a dialysis tubing (12-14 kDa MWCO) and dialyzed against DI water for 4 days, while replacing the DI water twice daily. The dialyzed solution was then transferred into 50-mL centrifuge tubes, snap-frozen in liquid nitrogen, and freeze-dried to obtain HA-MA, HA-Tau-20-MA, or HA-Tau-40-MA powder. The extent of methacrylation was determined using ^1^H-NMR.

### HA, HA-Tau-20, and HA-Tau-40 Nanosponge Synthesis

HA-MA, HA-Tau-20-MA, or HA-Tau-40-MA (60 mg) was dissolved in 800 µL of DI water, and 75 µL of lithium phenyl-2,4,6-trimethylbenzoylphosphinate (LAP; 4% w/v in DI water) was added. The solution was vortexed for 10 seconds and degassed in a bath sonicator for 10 minutes. The aqueous phase was then added dropwise to 10 mL of mineral oil containing 10% w/v ABIL EM 90 surfactant while stirring at 200 rpm. The water-in-oil solution was emulsified using a probe sonicator at 80% amplitude for 150 seconds (two 45-second pulses followed by two 30-second pulses). The resulting nanoemulsion was crosslinked via UV irradiation for 15 minutes under constant stirring at 300 rpm. The crosslinked nanoemulsion was poured into a chilled 1:1 acetone: hexane mixture, 15 times the volume of the mixture. The nanocarriers were pelleted by centrifugation at 7000 g for 10 minutes, and the supernatant was discarded. The pellet was washed three times with the 1:1 acetone: hexane mixture. The pellet was then dispersed in a 1:1 ethanol: water solution, transferred to a 12-14 kDa MWCO dialysis tube, and dialyzed against DI water for 3 days. The solution was subsequently transferred into a 50 mL centrifuge tubes, snap-frozen in liquid nitrogen, and freeze-dried to obtain the HA, HA-Tau-20, or HA-Tau-40 nanosponge powder.

### Characterization of Nanosponges

The nanosponges were characterized using dynamic light scattering (DLS) and zeta potential measurement. The freeze-dried nanosponges were redispersed in distilled water at a concentration of 100 μg/mL using gentle vortexing and sonication. The suspension was transferred into a disposable folded capillary cell (Cat. no. DTS1070; Malvern Panalytical), and the size and surface charge were measured using a Malvern Zetasizer Ultra-Red. The hydrodynamic diameter was determined by averaging three independent measurements, each consisting of 20 runs. The zeta potential was calculated by averaging three independent measurements, each consisting of 100 runs, and the Smoluchowski equation was used to determine the zeta potential value.

### *In Vitro* Protein Binding

The protein binding efficacy of the nanosponges was assessed using albumin and fibrinogen, the two most abundant plasma proteins. Bovine serum albumin (BSA; B-3371-50GM, AG Scientific) or fibrinogen (F8630, Sigma), at 10 µg/mL, was incubated with nanosponges (2.5 mg/mL) or PBS for 6 h at 37 °C with shaking at 70 rpm. The incubated solutions were centrifuged at 15,000 g for 20 minutes at 4 °C. The supernatant was carefully collected, incubated with Bradford dye (#5000006, Bio-Rad) for 10 minutes in a 96-well plate, and the absorbance was measured using an Infinite M200 multiplate reader (Tecan). The percentage change in protein concentration in supernatant relative to the total protein incubated (10 µg/mL) was plotted.

### Animal care

All animal procedures were conducted under protocol A116-23-05, approved by the Institutional Animal Care and Use Committee at Duke University, and were carried out in accordance with NIH guidelines and relevant national and international standards for laboratory animal care.

### *Ex Vivo* Plasma Cytokine Binding

The cytokine-scavenging efficacy of the nanosponge in plasma was assessed under *ex vivo* conditions. The plasma samples were collected 6 h after tibial fracture from aged C57BL/6J mice. Nanosponges in PBS were mixed with an equal volume of plasma to obtain a final nanosponge concentration of 2.5 mg/mL, then incubated for 6 h at 37 °C with orbital shaking at 70 rpm. Plasma samples treated with an equal volume of PBS served as the control. The incubated solutions were centrifuged at 15,000 g for 20 minutes at 4 °C. The supernatant was carefully collected without disturbing the pellet and analyzed for cytokine concentrations by ELISA according to the manufacturer’s protocol. ELISAs were performed for IL-1β (#ELM-IL1b-1, RayBiotech), IL-6 (#ELM-IL6-1, RayBiotech), and TNF-α (#ELM-TNFα-1, RayBiotech). For each cytokine, the percentage change in cytokine concentration in the nanosponge-treated supernatant relative to the PBS control was calculated and plotted.

### Plasma Sample Preparation and Proteomic Analysis

The freeze-dried nanosponge was dispersed in PBS at 10 mg/mL and allowed to hydrate for 6 h or overnight. The suspension was ultrasonicated on ice for 30 seconds (three times for 10 seconds each) then bath sonicated for 1 minute. 250 µL of the HA-Tau nanosponge was added to the 750 μL of mouse or human plasma (**Supplementary Table 4)** and mixed thoroughly. The final concentration of HA-Tau-40 nanosponge in the mixture was 2.5 mg/mL. The mixture was incubated overnight at 37 °C under shaking (75 rpm). The suspension was centrifuged for 20 minutes at 17000 g at 4 °C. The supernatant was carefully removed, and the nanosponge pellet was collected. 200 μL of ice-cold PBS was added to the pellet and mixed thoroughly. The suspension was centrifuged for 20 minutes at 17000 g at 4 °C. The supernatant was removed, and the washing step was repeated. Finally, the pellet was treated with 200 μL of the protein dissolution buffer (5% SDS, 25 mM Tris, 10 mM DTT, 50 mM NaCl, pH=8). The sample was heated at 70 °C in a water bath for 10 minutes. The resulting solution was further centrifuged at 25000 g for 20 minutes at 4 °C, and the supernatant was collected. The resulting protein extract was spiked with 1 pmol of bovine casein (Sigma) as an internal digestion control. Samples were then reduced for 15 min at 80 °C with 10 mM dithiolthreitol and alkylated with 20 mM iodoacetamide for 30 min at room temperature. Samples were then supplemented with a final concentration of 1.2% phosphoric acid and 375 µL of S-Trap (Protifi) binding buffer (90% MeOH/100mM TEAB). Proteins were trapped on the S-Trap micro cartridge, digested using 20 ng/µL sequencing grade trypsin (Promega) for 1 hr at 47 °C, and eluted using 50 mM TEAB, followed by 0.2% FA, and lastly using 50% ACN/0.2% FA. All samples were then lyophilized to dryness. Samples were resuspended in 40uL of 1% TFA/2% acetonitrile with 12.5 fmol/µL of yeast ADH. LC/MS/MS was performed using an EvoSep One UPLC coupled to a Thermo Orbitrap Astral high resolution accurate mass tandem mass spectrometer (Thermo). Briefly, 10% (approximately 300 ng) of each sample loaded onto an EvoTip was eluted onto a 1.5 µm EvoSep 150um ID x 15cm performance (EvoSep) column using the SPD30 gradient at 45 °C. Data collection on the Orbitrap Astral mass spectrometer was performed in a data-independent acquisition (DIA) mode of acquisition with a r=240,000 (@ m/z 200) full MS scan from m/z 380-980 in the OT with a target AGC value of 4e5 ions. Fixed DIA windows of 4 m/z from m/z 380 to 980 DIA MS/MS scans were acquired in the Astral with a target AGC value of 5e4 and max fill time of 6 ms. HCD collision energy setting of 27% was used for all MS2 scans. Data was imported into Spectronaut 19 (Biognosis). Relative peptide abundance was measured based on MS2 fragment ions of selected ion chromatograms of the retention time aligned runs. The MS/MS data was searched against a SwissProt Homo sapien or Mus musculus database (downloaded in 2024), a common contaminant/spiked protein database (bovine albumin, bovine casein, yeast ADH, etc.), and an equal number of reversed-sequence “decoys” for false discovery rate determination. A Direct DIA+ library free approach within Spectonaut was used to perform database searches. Database search parameters included fixed modification on Cys (carbamidomethyl) with variable modification on Met (oxidation). Full trypsin enzyme rules were used along with 10 ppm mass tolerances on precursor ions and 20 ppm on product ions. Spectral annotation was set at a maximum 1% peptide false discovery rate based on q-value calculations. Note that peptide homology was addressed using razor rules in which a peptide matched to multiple different proteins was exclusively assigned to the protein that has more identified peptides. Protein homology was addressed by grouping proteins that had the same set of peptides to account for their identification. A master protein within a group was assigned based on % coverage. Following protein identification and grouping, comparative and pathway enrichment analyses were performed to characterize the biological significance of the nanosponge-bound mouse and human proteome. The number of identified proteins per human sample was compared. List comparison was conducted in base R to identify unique proteins per sample [HR1.1] and illustrated as a Venn Diagram. Data was analyzed using Ingenuity Pathway Analysis (IPA) (QIAGEN Inc.; accessed on 02/18/2026). The User Dataset was set as the reference. The top 300 expressed proteins were used as input [HR2.1]. Analysis was conducted to identify canonical pathways, upstream regulators, and associated networks. Pathways were considered significantly enriched if the Benjamini-Hochberg-adjusted p value was less than 0.05. Network analysis was performed for the top enriched canonical pathway and its upstream regulators (p value < 0.05). Network diagrams were generated with nodes colored by expression intensity and predicted regulators were positioned at the center.

### *In Vivo* Biodistribution of Nanosponge

Biodistribution studies were performed in aged C57BL/6J mice (20-22 months). Each mouse received an intraperitoneal injection of 200 μL of Cy5-conjugated HA-Tau-40 nanosponge (150 mg/kg). Blood samples were collected at predetermined time points (0 min, 30 min, 1 h, 3 h, 6 h, 12 h, and 24 h) in heparin-coated tubes. A 20 μL aliquot of blood was diluted with 180 μL of PBS in a 96-well plate, and Cy5 fluorescence was measured using an Infinite M200 multiplate reader (Tecan) at an excitation wavelength of 615 nm and emission at 680 nm. A standard curve correlating HA-Tau-40-Cy5 nanosponge concentration with Cy5 fluorescence intensity was used to quantify the nanosponge concentration in blood samples. The total blood volume of each mouse was assumed to be 2 mL. The percentage of nanosponge remaining in circulation at each time point, relative to the total amount injected (25 mg/ml), was plotted as a function of time. After 24 h of nanosponge injection, the mice were euthanized, and all major organs, including the heart, liver, brain, lungs, spleen, and kidneys, were collected and fixed with 4% paraformaldehyde. The organ weights were recorded, and Cy5 fluorescence was imaged with the IVIS system. Fluorescence intensity, after subtracting background, was normalized to organ weight. Data analysis was performed using Living Image® software version 4.8.2 (Revvity), and the results were expressed as radiant efficiency per gram of tissue.

### Tibial fracture and nanosponge treatment

Tibial fractures in aged C57BL/6J mice (20—22 months) were induced as described previously [29]. Briefly, animals were anesthetized with 2% isoflurane and administered buprenorphine (1 mg/kg, sustained release; Zoopharm) for analgesia. A small incision was made to expose the right knee joint, after which a 0.7 mm pin was inserted from the tibial plateau through the medullary cavity to stabilize the bone. The surrounding muscle was carefully detached from the tibial surface, and the periosteum was stripped. A midshaft fracture was then created using surgical scissors. The wound was sutured, and two drops of bupivacaine (0.5%, Hospira) were applied topically along the incision site before animals were allowed to recover. Thirty minutes after fracture induction, mice received an intraperitoneal injection of either saline or HA-Tau-40 nanosponges (150 mg/kg).

### Blood collection and cytokine analysis

Peripheral blood was collected from the tail vein of mice 6 h after surgery. Samples were drawn into anticoagulant-coated tubes and centrifuged at 2000 × g for 15 min at 4 °C to separate plasma. The resulting plasma was stored at −80 °C until analysis. Cytokine levels of IL-1β (#ELM-IL1b-1, RayBiotech), IL-6 (#ELM-IL6-1, RayBiotech), and TNF-(#ELM-TNFα-1, RayBiotech) were measured using ELISA kits following the manufacturers’ instructions.

### Immunohistochemical analysis

Immunostaining of brain tissue sections was performed using standard free-floating protocols. Sections were washed three times with PBS (5 min each) and permeabilized in 0.1% Triton X-100 for 15 min. After blocking with 3% BSA in PBS for 1 h at room temperature, sections were incubated overnight at 4 °C with primary antibodies (**Supplementary Table 1**). Following three washes, sections were incubated with the appropriate secondary antibodies (**Supplementary Table 2**) for 1 h at room temperature. Stained sections were mounted on glass slides using DAPI-containing mounting medium and imaged using either a Keyence BZ-X700 microscope or a Dragonfly Spinning Disk Confocal Microscope (Andor). Fluorescence thresholds for each channel were fixed based on both negative and positive controls. Image analysis was performed in ImageJ (version 1.53i). Images were converted to 8-bit grayscale, thresholded, and quantified using the “Measure” tool. Fluorescence intensity values represent the average from at least two sections per animal, with a minimum of five animals (biological replicates) per group.

### Synaptophysin/PSD95 imaging and colocalization analysis

For quantitative analysis of synaptic puncta, brain sections were immunostained for Synaptophysin and PSD95 and subsequently imaged using a Dragonfly Spinning Disk Confocal Microscope (Andor) with a 63× oil-immersion. Z-stack images spanning 5 μm in thickness were acquired from the middle portion of the tissue to minimize potential staining artifacts using a 0.3 μm step size. To account for regional differences in synaptic puncta density, images were acquired from anatomically matched regions (CA1) using the same imaging parameters across all experimental groups. For image preprocessing, brightness and contrast were adjusted individually for each image to account for variations in background fluorescence, while minimizing background signal and preserving clearly identifiable synaptic puncta. Synaptophysin and PSD95 puncta were quantified using the SynBot plugin in ImageJ [50]. For quantitative analysis, the same predefined analysis parameters, including threshold settings, were applied uniformly to all images across experimental groups. The percentage of colocalized puncta was calculated as the number of colocalized puncta divided by the total number of puncta detected in both channels (Synaptophysin + PSD95) × 100. At least two sections were analyzed per animal, and the mean value from each animal was used as the biological replicate for statistical analysis.

### Gene expression analysis

Total RNA was extracted from mouse hippocampal tissue using Trizol reagent. Briefly, tissue was homogenized in 1.5 mL of Trizol and centrifuged at 12,000 × g for 20 min at 4 °C. The supernatant was transferred to a new tube, mixed with 300 μL chloroform, and centrifuged at 12,000 × g for 20 min at 4 °C. The aqueous phase was collected, combined with 1 mL of 100% ethanol, and centrifuged at 10,000 × g for 10 min at 4 °C. The resulting RNA pellet was washed, air-dried, and eluted in 50 μL of RNase-free water. One microgram of total RNA was reverse transcribed into cDNA using the iScript cDNA Synthesis Kit (1708891, Bio-Rad) following the manufacturer’s protocol. Quantitative real-time PCR (qRT-PCR) was performed using the Bio-Rad CFX96 Touch cycler under the following conditions: initial denaturation at 95 °C for 30 s (1 cycle), followed by 40 cycles of 95 °C for 5 s and 60 °C for 30 s. Primer sequences are listed in **Supplementary Table 3**. Gene expression levels were normalized to housekeeping gene expression and analyzed relative to the corresponding controls, with results presented as fold change.

### BBB permeability analysis using Evans blue dye

Blood–brain barrier (BBB) permeability was assessed using Evans blue dye as previously described [51]. Briefly, 100 µL of 4% Evans blue solution prepared in 0.9% saline was injected intravenously 23 h post-fracture. One h after injection, mice were transcardially perfused with ice-cold PBS followed by 4% paraformaldehyde (PFA). Brains were collected, post-fixed, and processed for cryosectioning. The sections were washed three times with PBS and immunostained with the endothelial cell marker CD31. Stained sections were imaged using a Keyence fluorescence microscope, and the area of extravascular Evans blue dye was quantified using ImageJ software.

### *Ex vivo* explant culture

Brains were collected from 22-month-old male mice and washed with cold PBS. Coronal brain slices (∼1 mm thick) containing the hippocampal regions were prepared using a rodent brain matrix (RBM-2000C, ASI Instruments) according to the manufacturer’s instructions. Slices were immediately transferred to brain explant culture medium consisting of Neurobasal medium supplemented with B27, N2, GlutaMAX, penicillin–streptomycin, and horse serum, and incubated at 37 °C in 5% CO and 95% humidity for 2 h. Brain slices were then cultured for 6 h in explant media containing either 5% post-fracture plasma from 22-month-old mice or 5% nanosponge-conditioned post-fracture plasma (n = 3). Slices maintained in regular explant media served as controls. Following treatment, one half of the slices were fixed in 4% paraformaldehyde, cryo-embedded, sectioned, and immunostained for glial activation markers IBA1 and GFAP. Sections were imaged using a Keyence fluorescence microscope, and fluorescence intensity was quantified with ImageJ. Hippocampal regions from the other half of the slices were punched and total RNA was extracted using TRIzol reagent as described above. Quantitative PCR was performed to determine the expression of inflammatory markers (Il-6, Il1β, and Tnf-α). Expression levels were normalized to the housekeeping gene and expressed relative to their respective controls. Primer sequences are provided in Supplementary **Supplementary Table 3**.

### MEA analysis

Mouse hippocampal explants were carefully positioned onto a 32-electrode multi-electrode array (MEA; Harvard Biosciences #890385). The MEA was connected to a CerePlex Direct data acquisition system via a digital headstage interface (Blackrock Neurotech, # 11259). Prior to recording, explants were allowed to equilibrate for 5 minutes on the electrode to ensure optimal recording conditions. Spontaneous extracellular electrical activity was recorded for 5 minutes at a sampling rate of 30 kS/s. Raw signals were band-pass filtered using a 4-pole Bessel filter with a 300 Hz low-cutoff to isolate spiking activity. Spike detection and sorting were performed offline using Offline Sorter software (Plexon, # 99952-402). Waveforms were clustered using the valley-seeking method, with a sigma parameter ranging from 4 to 6 to optimize the separation of neuronal units. Only units with a signal-to-noise ratio above 2 and consistent waveform morphology across the recording were included in subsequent analyses.

### Y-maze analysis

Spatial working memory was assessed using the Y-maze spontaneous alternation task. The maze consists of three identical arms positioned at 120° angles (Maze Engineers). Mice were placed at the end of one arm and allowed to freely explore all three arms for 8 minutes. The sequence of arm entries was recorded and analyzed manually, and distance tracking and visualization was conducted using ezTrack.[52] Spontaneous alternation behavior was calculated as the percentage of successive entries into three different arms on overlapping triplet sets, using the formula: Spontaneous alternation (%) = (Number of alternations) / (Total arm entries−2) ×100.

### *In vitro* human BBB transwell model

Primary Human Brain Microvascular Endothelial Cells (# ACBRI 376, Cell Systems) monolayers were cultured on polyester Transwell inserts (# 3470, Costar). Wells were coated with a chilled PBS–Matrigel mixture to enhance cell adhesion. After incubation for 1 h at 37 °C, excess coating solution was removed, and wells were washed twice with PBS. Endothelial cells were seeded at a density of 26,000 cells/well in 100 µL of EGM-2MV medium (#CC-3202, Lonza) added to the apical chamber, while 500 µL of media was added to the basolateral chamber. Cells were cultured until a confluent monolayer was established. For preparation of nanosponge-conditioned human plasma, 250 µL of HA-Tau-40 nanosponge (10 mg/mL) was added to 750 µL of human plasma and mixed thoroughly. The mixture was incubated overnight at 37 °C with shaking at 75 rpm and subsequently centrifuged at 17,000 × g for 20 min at 4 °C. The resulting supernatant was carefully collected and used as nanosponge-conditioned human plasma for experiments. Confluent endothelial monolayers were exposed for 6 h to either regular culture medium, medium containing 5% aged human plasma, or medium containing 5% nanosponge-conditioned aged human plasma. Transendothelial electrical resistance (TEER) was subsequently measured using a custom-built Arduino-based circuit that generated a 5 V, 12.5 Hz square wave (half-period = 40 ms) across the Transwell. Voltage (mV) and current (µA) were measured simultaneously using two AC multimeters connected in parallel and series, respectively. Measurements were recorded every 30 seconds for 10 minutes using a camera to ensure consistency. The resistance (R) across the monolayer was calculated from Ohm’s law (R = V/I), and the resistance attributable to the cell layer was determined by subtracting the resistance of blank wells. The final TEER (Ω · cm²) was calculated as:

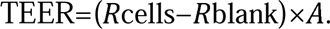

where *A* =0.3318 cm^2^ is the membrane area.

BBB permeability was quantified using fluorescein isothiocyanate (FITC)-labeled dextran (10 kDa) as a fluorescent tracer. Prior to tracer introduction, media was removed from the apical (top) chamber. FITC–dextran (10 ug/mL; 100 μL) was added to the apical chamber. At 60 min time point, aliquots were collected from the basolateral chamber to quantify dextran diffusion across the endothelial monolayer. Fluorescence intensity was measured using a microplate reader (excitation: 485 nm, emission: 528 nm) and the concentration of transported FITC–dextran was determined using a standard curve generated from known FITC–dextran concentrations.

### H and E staining

Liver tissues were harvested 24 h after nanosponge or saline administration and fixed in 4% paraformaldehyde (PFA). Following fixation, tissues were cryoprotected in 30% sucrose, embedded in optimal cutting temperature (OCT) compound, and sectioned at a thickness of 15 μm using a cryostat. Sections were mounted on glass slides and washed twice with distilled water prior to staining. For hematoxylin and eosin (H&E) staining, sections were first stained with Mayer’s hematoxylin (# 26043-06, Electron Microscopy Sciences) for 3 min to visualize nuclei. Slides were then rinsed with multiple changes of distilled water until excess dye was removed. Nuclear bluing was achieved by incubating the sections in 1× PBS for 3 min, followed by washing with three changes of distilled water. Sections were briefly rinsed in 95% ethanol for 1 min and subsequently counterstained with alcoholic eosin (# 26051-21, Electron Microscopy Sciences) for 1 min to visualize cytoplasmic structures. Following staining, sections were dehydrated through three sequential washes in 95% ethanol and three washes in 100% ethanol (1 min each). Slides were then cleared in xylene (three changes, 1 min each), mounted with mounting medium, and coverslipped. Stained sections were imaged using imaged using a Keyence (BZ-X710) microscope.

### Statistical analysis

Statistical analyses were performed using GraphPad Prism 10.2.1. Comparisons between two independent groups were performed using a two-tailed unpaired *t*-test. For comparisons among three or more groups, one-way analysis of variance (ANOVA) followed by Tukey’s multiple-comparisons test was used. Each data point represents an individual animal (biological replicate). Data are presented as mean ± standard deviation (SD). A *p*-value < 0.05 was considered statistically significant.

## Data availability

All data generated or analyzed during this study are included in the manuscript.

## Acknowledgements

The authors thank Yuru Vernon Shih for assistance with tibial fracture surgery and Jiaul Hoque for assistance with intravenous (IV) injections. This work was funded by the National Institutes of Health (R01AR071552 and R01AR079189).

