## Supplementary Information for "Engineered nanosponges mitigate peripheral stress-induced neuroinflammation and restore cognitive function"

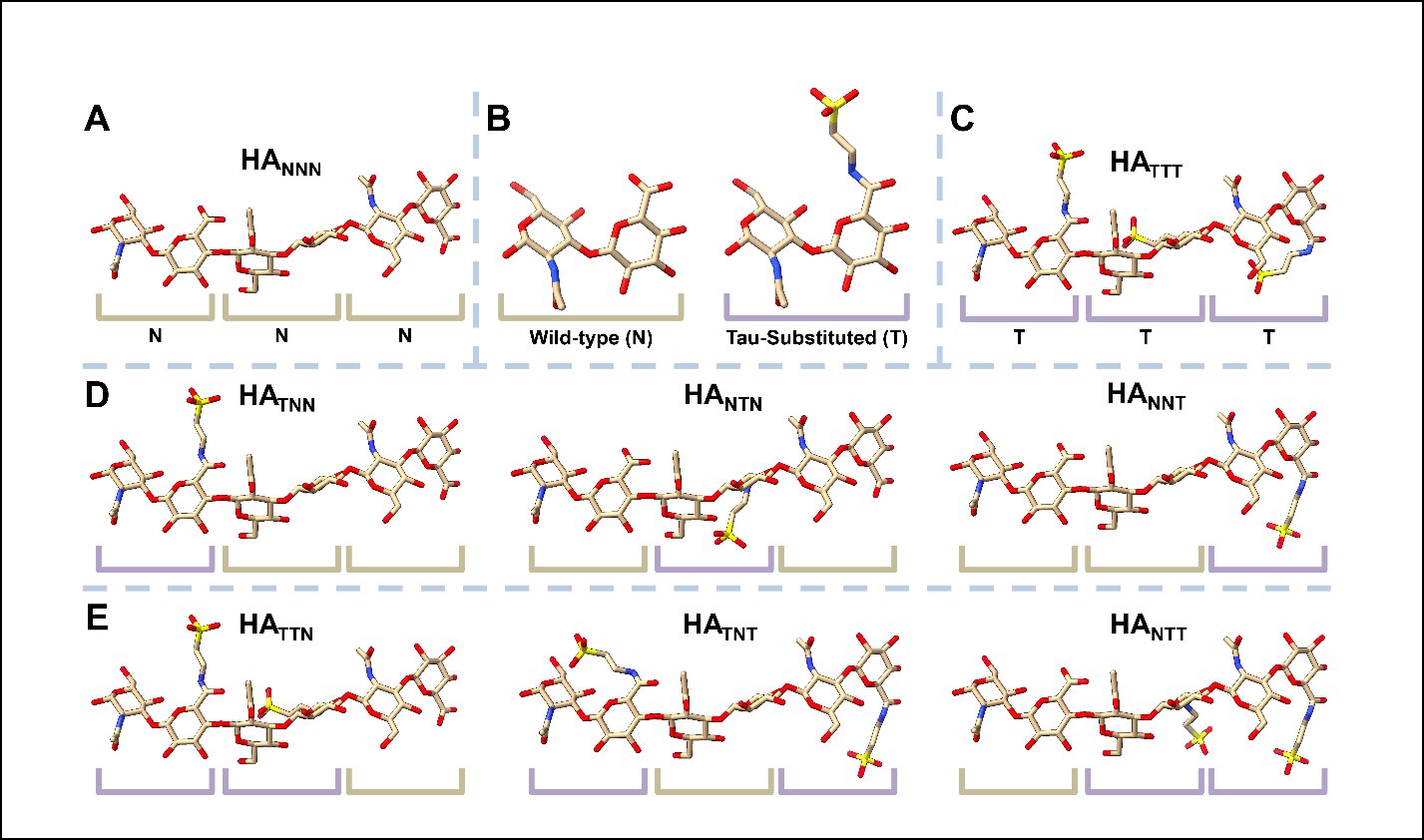

**Supplementary Fig. 1.** Hyaluronic acid trimeric oligomers examined in docking studies. A) Wild-type HA trimer. B) Unsubstituted (wild-type) HA monomeric unit labeled by the letter N with a yellow bracket and a Tau-substituted unit is labeled by the letter T with a gray bracket. C) Fully Tau-substituted HA trimer. D) 1/3 Tau-substituted trimers. E) 2/3 Tau-substituted trimers.

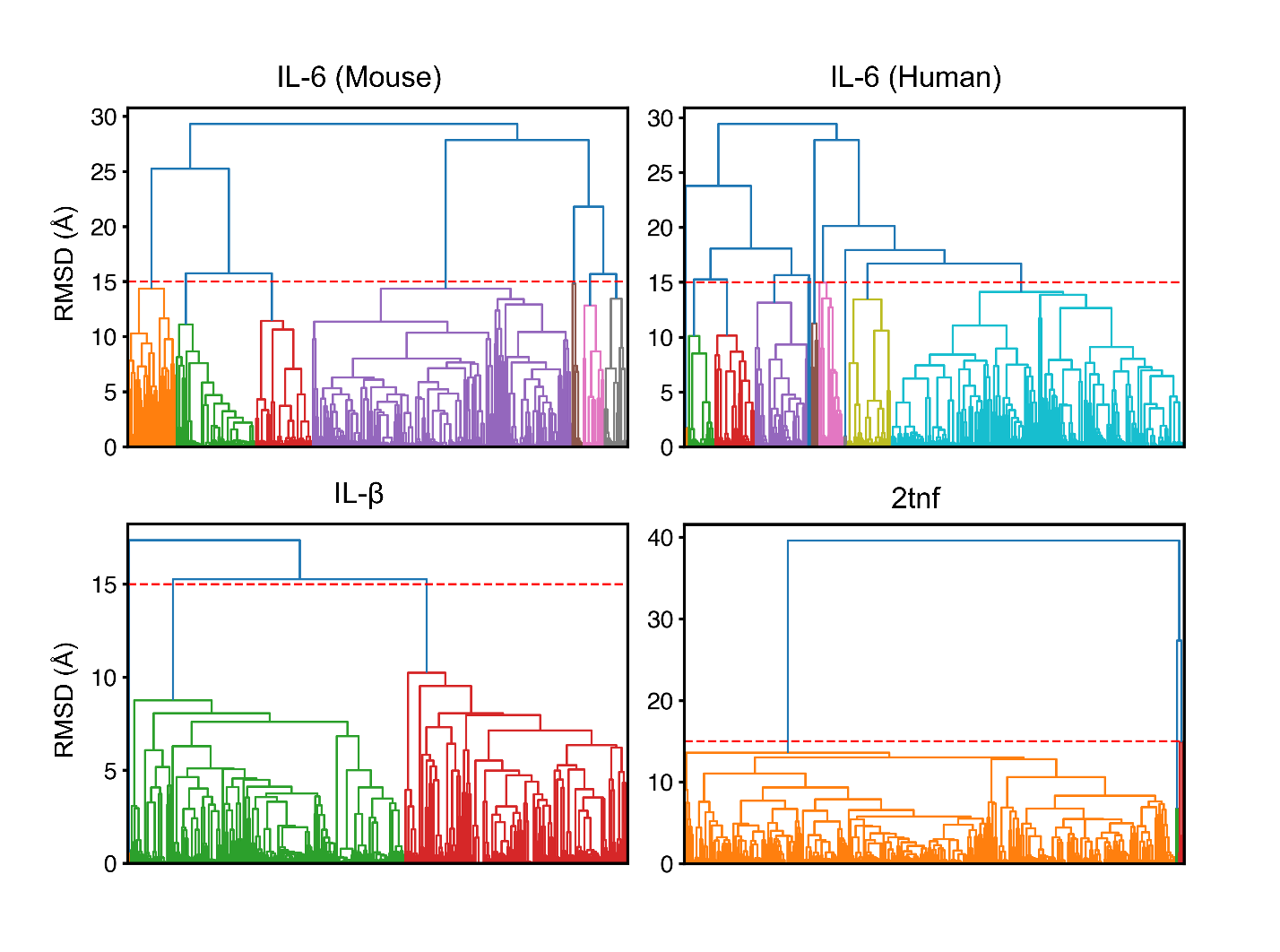
 **Supplementary Fig. 2.** Clustering of predicted docking conformations. Clusters were separated using a RMSD threshold of 15 angstrom (dotted red line) and grouped using the average linkage method (cluster is colored individually).

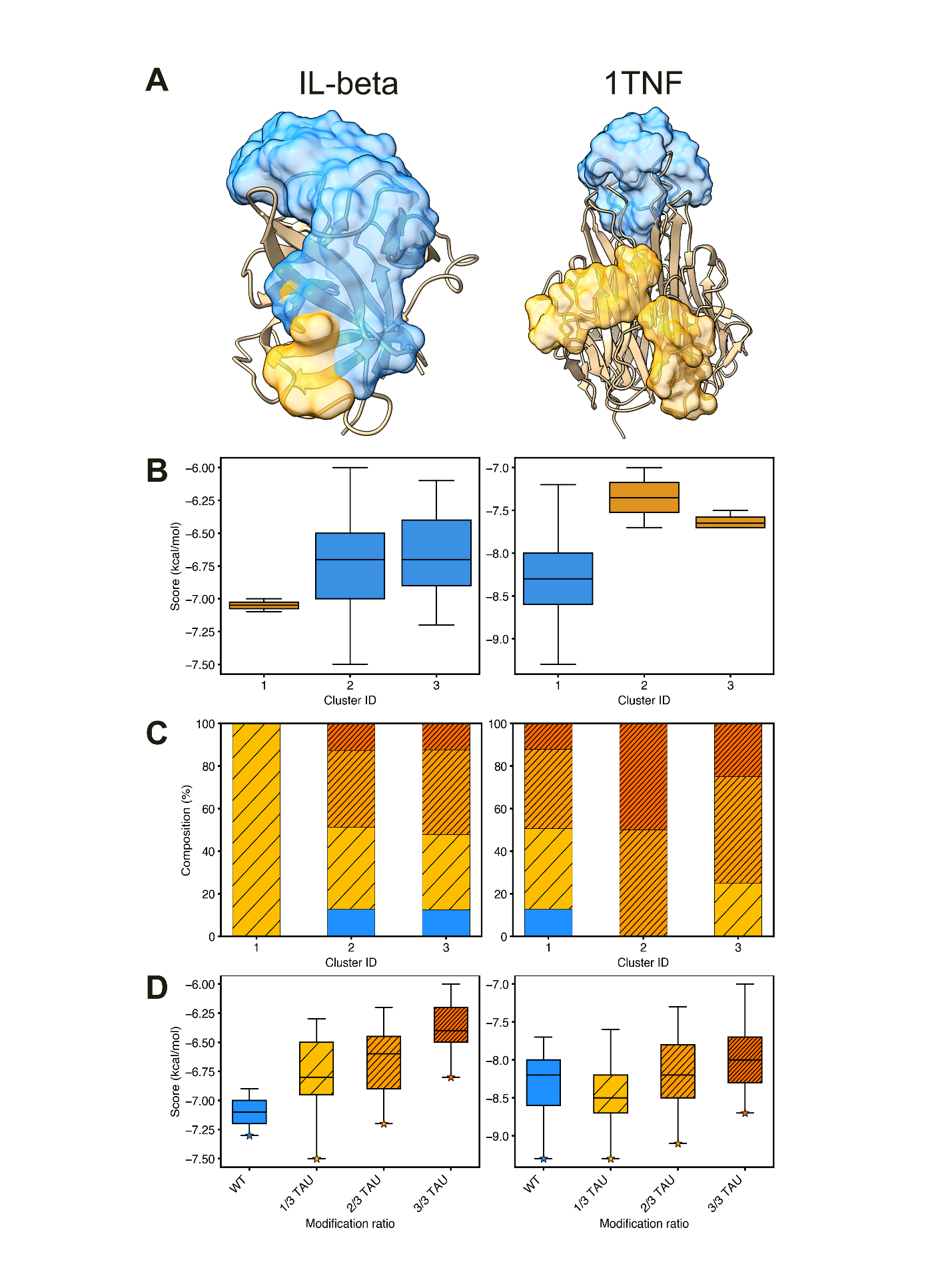

**Supplementary Fig. 3.** HA trimer docking results onto IL-beta (left) and 1TNF (right). A) Clusters visualized onto each IL-beta and 1TNF, colored based on if the cluster contains only unsubstituted HA (orange) or includes Tau-substituted HA (blue). B) Predicted docking scores grouped by cluster, again colored based on the cluster’s composition. C) Composition of each cluster: blue is unsubstituted HA (blue), 1/3 HA-Tau (yellow), 2/3 HA-Tau (orange), and fully substituted (dark orange). D) Predicted docking scores grouped by HA-Tau ratio, following the same color scheme as panel C. Stars indicate the best predicted binding score.

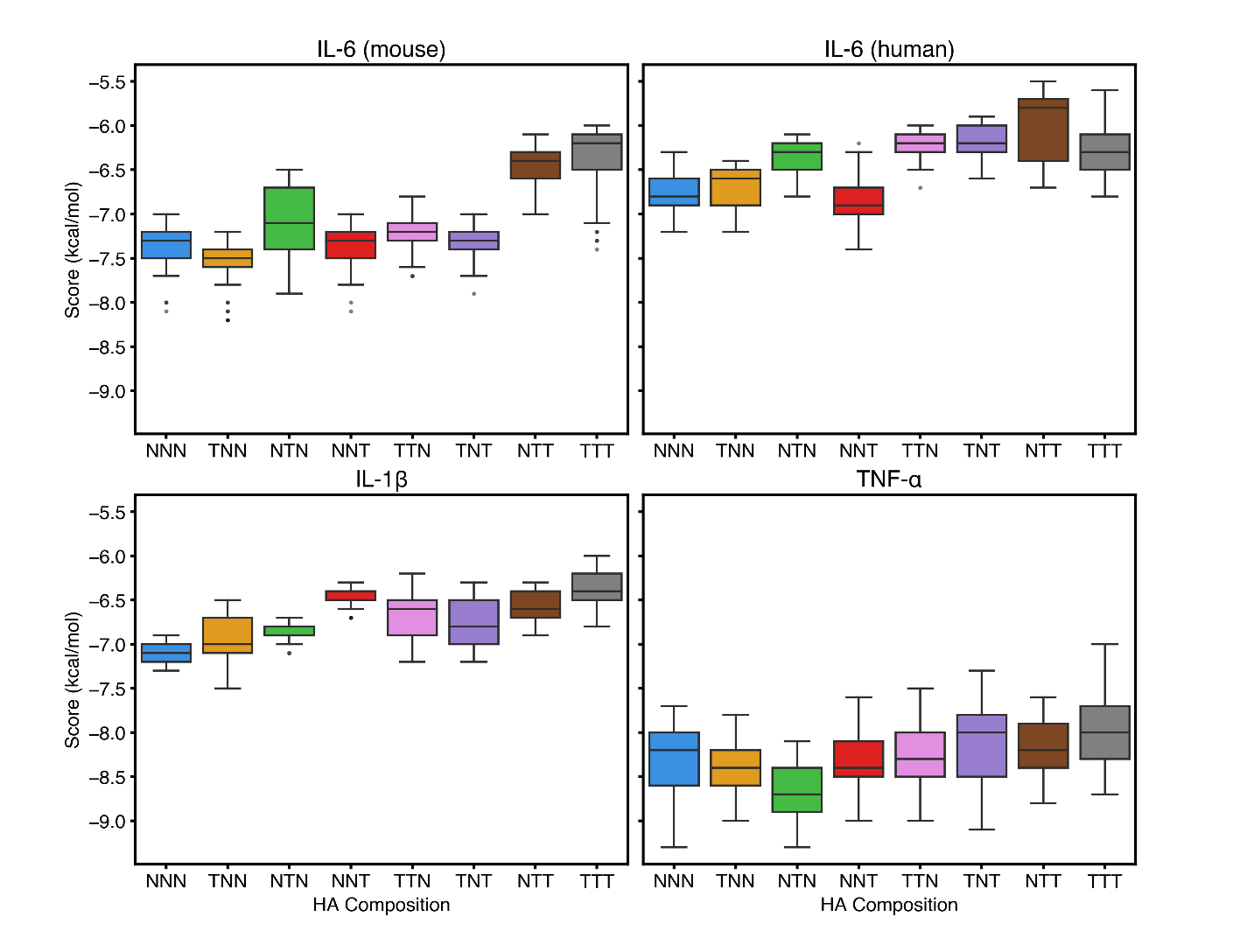

**Supplementary Fig. 4.** Predicted binding scores grouped by HA trimer variants.

**Supplementary Fig. 5**. ^1^H-NMR spectrum of the A) HA-Tau 20 and B) HA-Tau 40 polymer
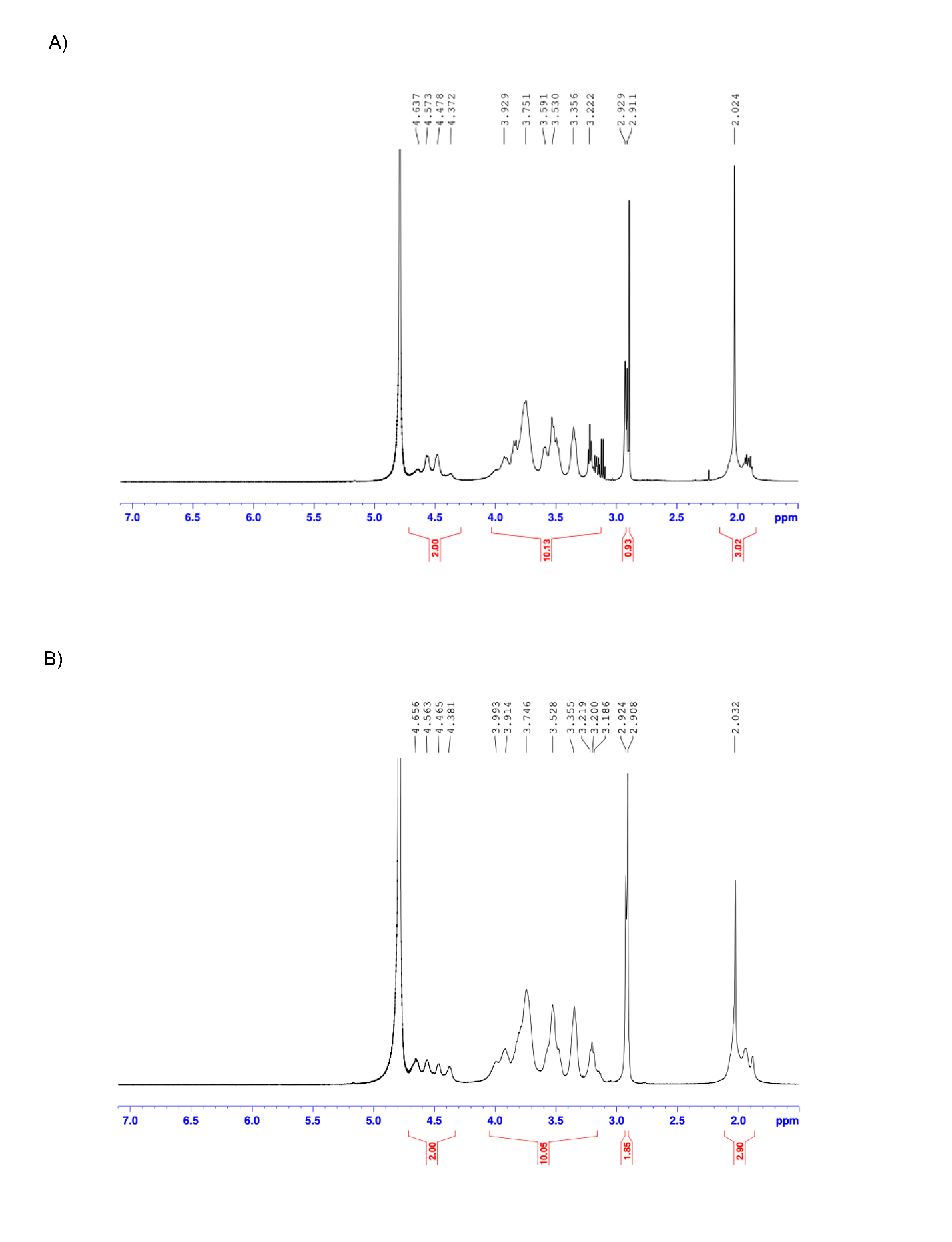
.

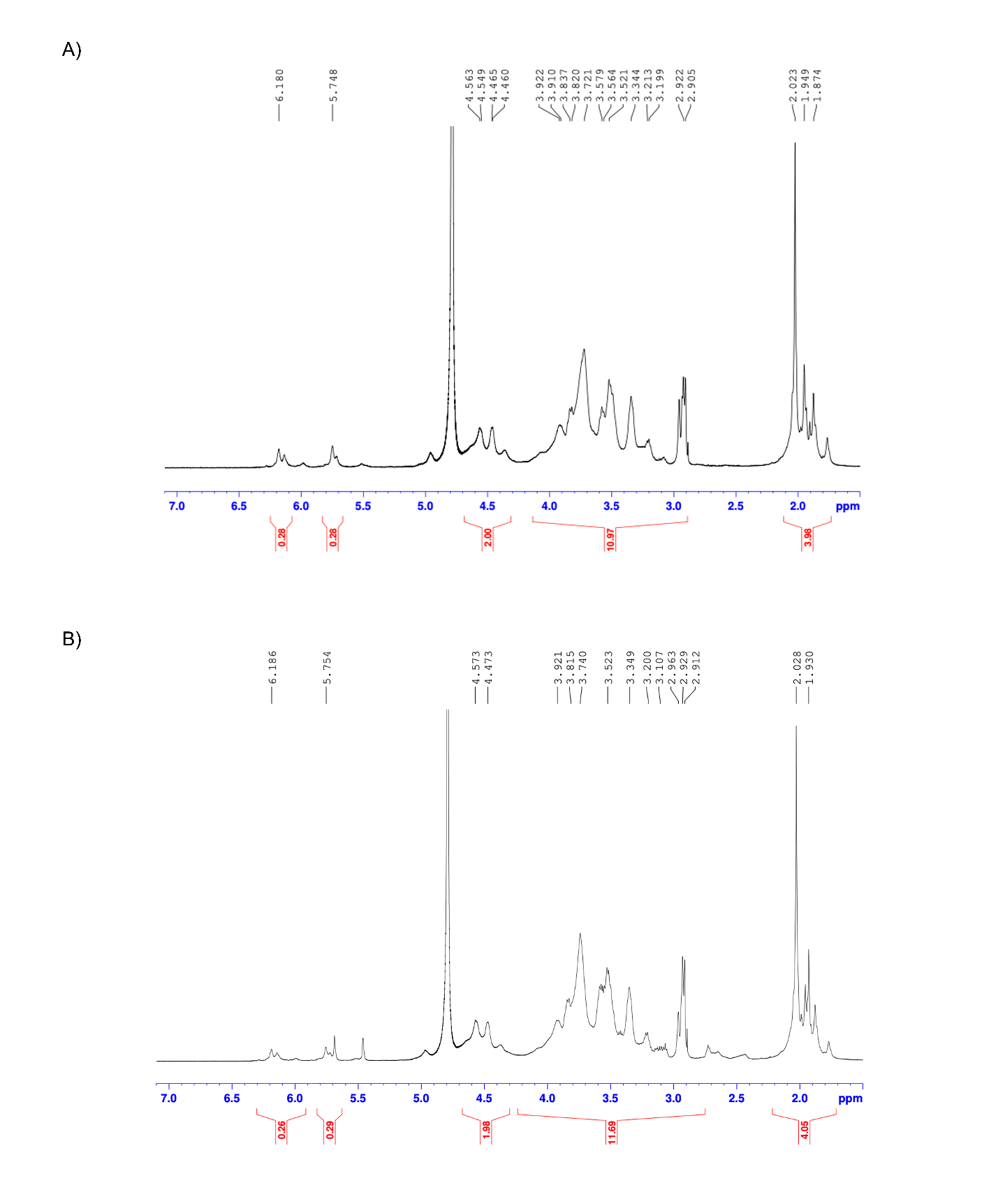

**Supplementary Fig. 6**. ^1^H-NMR spectrum of the HA-Tau-20-MA and HA-Tau-40-MA polymer.

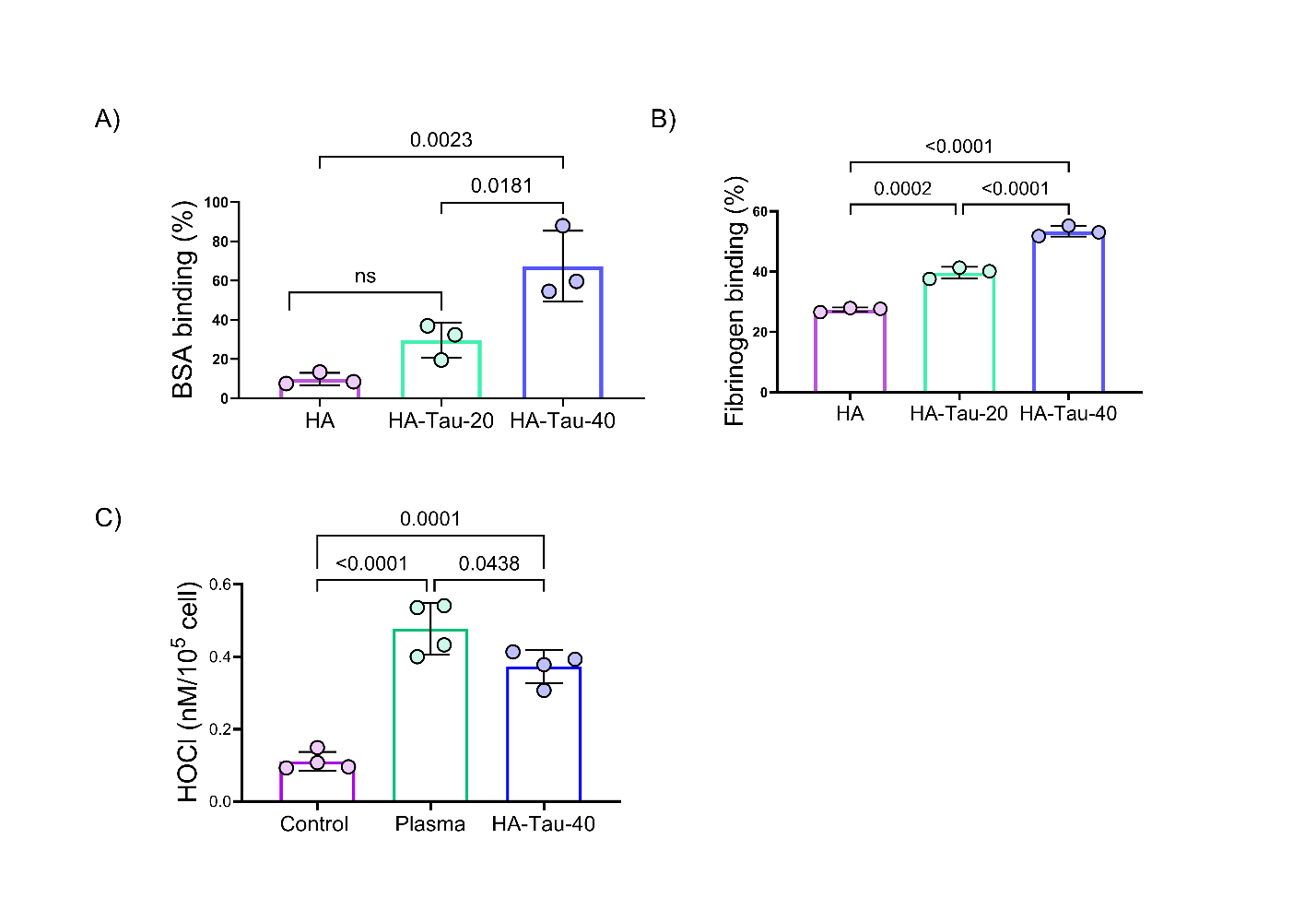

**Supplementary Fig. 7. In vitro protein binding and HOCl neutralization by HA–Tau nanosponges**. (A) BSA binding by HA, HA–Tau-20, and HA–Tau-40 nanosponges (n=3). (B) Fibrinogen binding by HA, HA–Tau-20, and HA–Tau-40 nanosponges (n=3). (C) HOCl levels in the culture supernatant of primary human neutrophils exposed to post-fracture plasma in the presence of HA–Tau-40 nanosponges (n=4). Data represent mean ± s.e.m. Statistical analyses were performed using one-way ANOVA with Tukey’s multiple comparisons test in GraphPad Prism 10.2.1. P values less than 0.05 are shown.

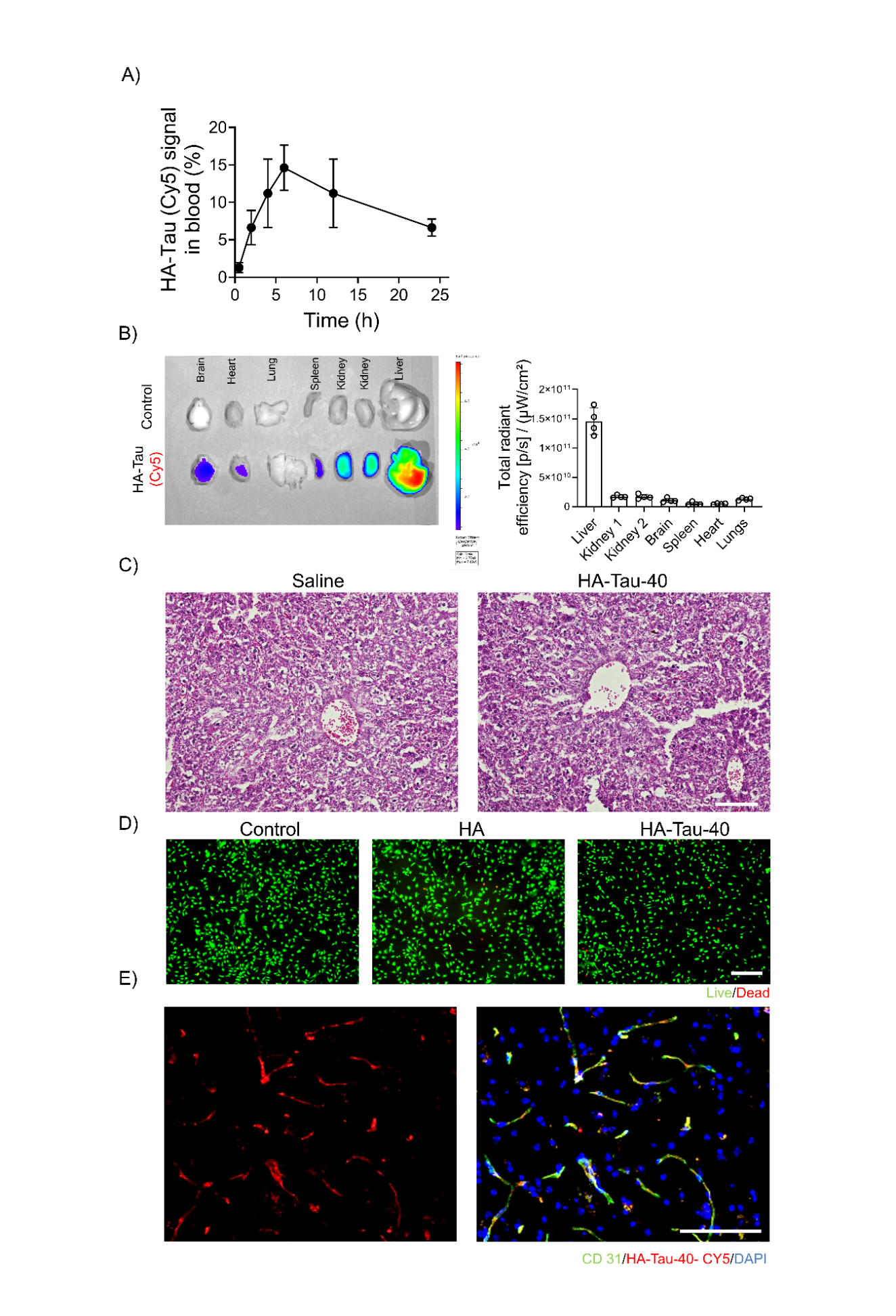

**Supplementary Fig. 8. *In vivo* biodistribution and cytotoxicity of HA–Tau nanosponges.** A) Longitudinal quantification of Cy5.5-labeled HA–Tau nanosponge fluorescence in blood following intraperitoneal administration in aged mice (n=3). B) IVIS images of major organs collected 24 h after administration of Cy5.5-labeled HA–Tau nanosponges. Quantification of fluorescence signals in individual organs is shown on the right (n=4). C) Representative hematoxylin and eosin (H&E)-stained liver sections from saline- and HA–Tau-40-treated mice 24 h after administration. Scale bar, 100 μm. D) Representative live/dead staining of primary mouse macrophages following 24 h exposure to HA, or HA–Tau-40 nanosponges and untreated controls. Scale bar, 100 μm. E) Representative fluorescence images of brain sections showing CD31-positive blood vessels (green), Cy5.5-labeled HA–Tau nanosponges (red), and nuclei (DAPI, blue). Scale bar, 100 μm.

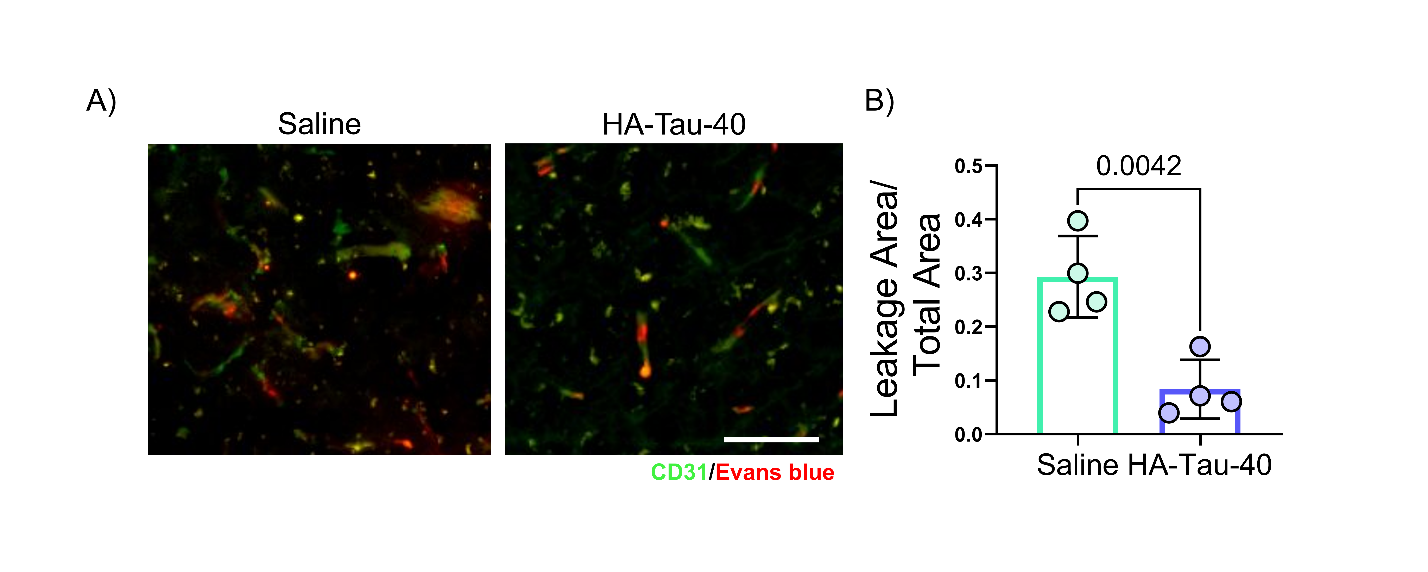

**Supplementary Fig. 9. HA–Tau nanosponges reduce blood–brain barrier permeability following tibial fracture.** A) Representative hippocampal images showing CD31-positive blood vessels (green) and Evans Blue fluorescence (red) in saline- and HA–Tau-40-treated mice following tibial fracture. Scale bar, 20 μm. B) Quantification of perivascular Evans Blue fluorescence normalized to the total tissue area is shown on the right (n=4). Statistical analyses were performed using two-tailed Student’s t-test in GraphPad Prism 10.2.1.

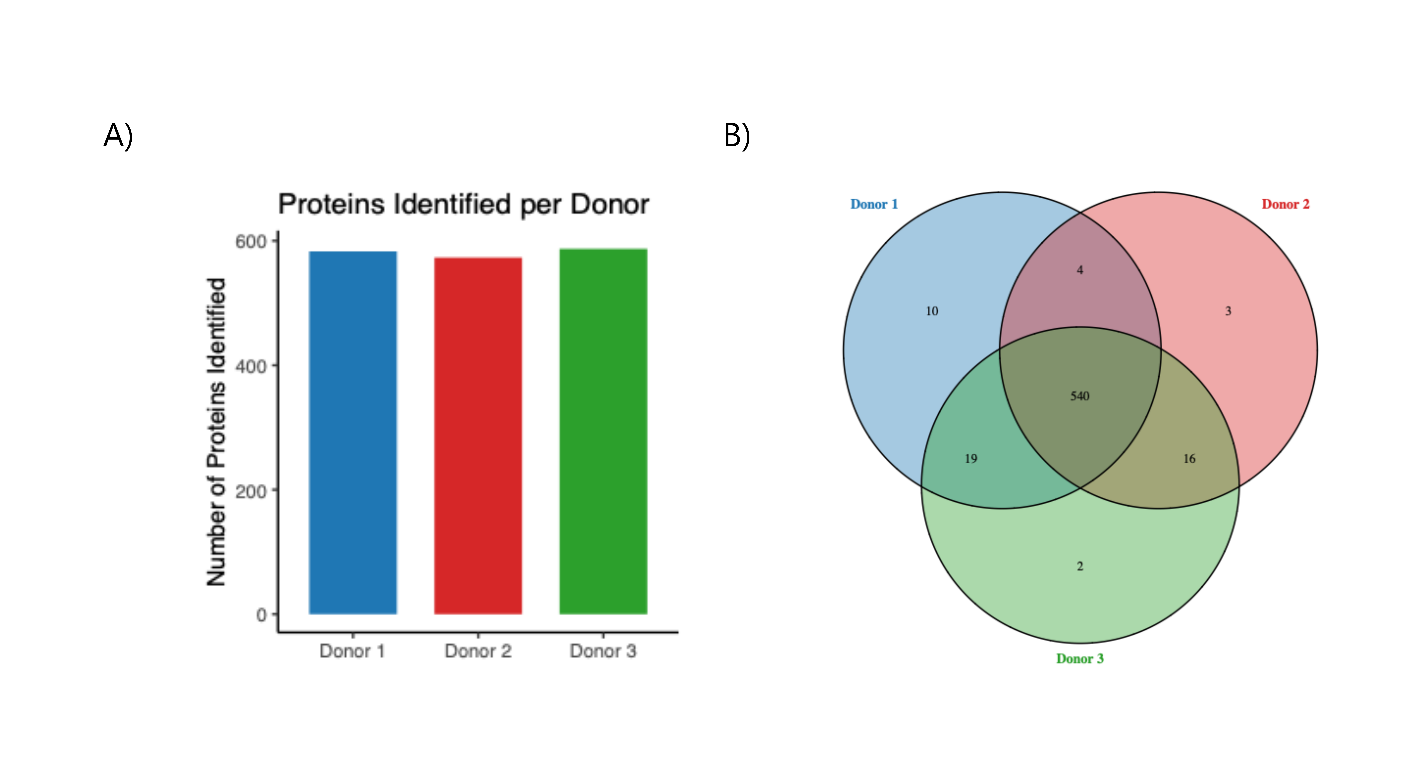
 **Supplementary Fig. 10. Proteomic profiling of human plasma proteins captured by HA–Tau-40 nanosponges.** A) Number of proteins identified in HA–Tau-40 nanosponge-associated protein coronas from plasma obtained from three independent aged human donors. B) Venn diagram showing the overlap of proteins identified across the three donors, with 540 proteins commonly identified among all donors.

**Supplementary Table 1.** List of primary antibodies.

| **Antibody** | **Company** | **Catalog#** | **Species** | **Dilution** |
| --- | --- | --- | --- | --- |
| Iba-1 | Waco | 011-27991 | Goat | 1:400 |
| CD68 | Bio-Rad | MCA1957GA | Rat | 1:400 |
| GFAP | Agilent Dako | Z033429-2 | Rabbit | 1:500 |
| Podocalyxin | R&D system | AF1556-SP | Goat | 1:300 |
| VCAM-1 | Invitrogen | MA5-11447 | Mouse | 1:100 |
| CD31/PECAM-1 | R&D Systems | AF3628-SP | Goat | 1:100 |
| Synaptophysin1 | Synaptic Systems | 101 011 | Mouse | 1:1000 |
| PSD-95 | Invitrogen | 51-6900 | Rabbit | 1:200 |

**Supplementary Table 2.** List of secondary antibodies

| **Antibody** | **Company** | **Catalog#** | **Species** | **Dilution** |
| --- | --- | --- | --- | --- |
| Alexa Fluor® 647 AffiniPure Donkey Anti-Rabbit IgG (H+L) | Jackson Immunoresearch | 711-605-152 | Donkey | 1:500 |
| Alexa Fluor® 488 AffiniPure Donkey Anti-Mouse IgG (H+L) | Jackson Immunoresearch | 715-545-150 | Donkey | 1:500 |
| Rhodamine Red™-X (RRX) AffiniPure Donkey Anti-Goat IgG (H+L) | Jackson Immunoresearch | 705-295-147 | Donkey | 1:400 |
| Alexa Fluor® 488 AffiniPure Donkey Anti-Rat IgG (H+L) | Jackson Immunoresearch | 712-545-153 | Donkey | 1:500 |
| Donkey anti-Mouse, Secondary Antibody, Alexa Fluor™ 647 | Thermofisher Scientific | A-31571 | Donkey | 1:500 |

**Supplementary Table 3.** List of primers used for RT-qPCR analysis

| m ZO1 (F) | 5'-GTTGGTACGGTGCCCTGAAAGA-3' |
| --- | --- |
| m ZO1 (R) | 5'-GCTGACAGGTAGGACAGACGAT-3' |
| m Glut1 (F) | 5'-GCTTCTCCAACTGGACCTCAAAC-3' |
| m Glut1 (R) | 5'-ACGAGGAGCACCGTGAAGATGA-3' |
| m Podxl (F) | 5'-ATAACCAGGCGGTGGCAGTGAA-3' |
| m Podxl (R) | 5'-CCAGCTTCATGTCACTGACTCC-3' |
| m Cldn5 (F) | 5'-TGACTGCCTTCCTGGACCACAA-3' |
| m Cldn5 (R) | 5'-CATACACCTTGCACTGCATGTGC-3' |
| m TNF-α (F) | 5'-AACTTCGGGGTGATCGGTCC-3' |
| m TNF-α (R) | 5'-TGGTTTGTGAGTGTGAGGGTCT-3' |
| m IL-1β (F) | 5'-TGCCACCTTTTGACAGTGATG-3' |
| m IL-1β (R) | 5'-ATGTGCTGCTGCGAGATTTG-3' |
| m IL-6 (F) | 5'-AAGACAAAGCCAGAGTCCTTCA-3' |
| m IL-6 (R) | 5'-GCATTGGAAATTGGGGTAGGAAG-3' |
| m 18S (F) | 5'-ACCAGAGCGAAAGCATTTGCCA-3' |
| m 18S (R) | 5'-ATCGCCAGTCGGCATCGTTTAT-3' |

**Supplementary Table 4**. Demographic characteristics of human plasma donors.

| Lot number | Gender | Race | Ethnicity | Age |
| --- | --- | --- | --- | --- |
| HMN143784 | M | White/caucasian | Hispanic/Latino | 75 |
| HMN143785 | M | Other | Hispanic/Latino | 71 |
| HMN143786 | M | White/caucasian | Hispanic/Latino | 73 |
